# Muscle Fiber- and Cell Type-Specificity of Training Adaptation in Male Mice

**DOI:** 10.1101/2025.11.04.686534

**Authors:** Sedat Dilbaz, Martina Baraldo, Peter Škrinjar, Volkan Adak, Regula Furrer, Alina Stein, Stefan A. Steurer, Gesa Santos, Christoph Handschin

**Author notes:** Authors contributed equally (Sedat Dilbaz, Martina Baraldo).

## Abstract

Skeletal muscle exhibits extraordinary structural and functional plasticity in response to contractile activity. As a syncytium within a complex microenvironment, its adaptation relies on the coordinated response of specialized myonuclear populations and diverse mononucleated cells. Here, we present a high-resolution single-nucleus RNA-sequencing atlas of 550’000 nuclei, capturing the longitudinal transcriptional dynamics of 17 myonuclear and 21 mononuclear populations following exhaustive exercise. We demonstrate that training status reshapes these cellular programs into divergent adaptive trajectories. While mononucleated cells serve as the primary mediators of intercellular communication during recovery, a subset of oxidative myonuclei enters a delayed regenerative state, reflecting a bifurcated response to the metabolic demands of endurance exercise. This is accompanied by signatures of impaired innervation and transient neuromuscular junction (NMJ) remodeling. Notably, prior training accelerates homeostatic recovery by preserving neuromuscular integrity and shielding oxidative nuclei from damage-associated signatures. Together, this atlas reveals that muscle plasticity emerges from a highly coordinated interplay of multicellular remodeling and provides a foundational resource for understanding the mechanisms of exercise adaptation.

## INTRODUCTION

Skeletal muscle is among the body’s most adaptable tissues, capable of extensive structural, metabolic, and functional remodeling in response to contractile activity. This plasticity underlies the systemic benefits of exercise, which protects against non-communicable diseases including cardiovascular decay, type 2 diabetes, neurodegeneration, and certain cancers^1^. However, the cellular mechanisms governing these adaptations remain poorly defined. While recent large-scale efforts have charted the multi-omic landscape of exercise^2–4^, including multi-omics studies from the Molecular Transducers of Physical Activity Consortium (MoTrPAC)^5–7^, these bulk approaches lack the resolution to reveal how specific fiber types and diverse muscle-resident mononucleated cells (MMCs) orchestrate tissue plasticity. Skeletal muscle is a complex syncytium where multinucleated fibers are embedded in a heterogeneous microenvironment of vascular, immune, stromal, and nerve-associated cells^8^. These populations occupy specialized functional niches, such as neuromuscular and myotendinous junctions, which likely serve as hubs for localized signaling^8–10^. Furthermore, the fiber-type spectrum, from slow twitch/oxidative/fatigue-resistant to fast twitch/glycolytic/fatigable, imposes distinct metabolic demands upon endurance- or strength-type of movement that likely dictate divergent adaptive responses^11^. In simplified terms, these fibers can be classified as type I, IIA, IIX, and, in rodents, type IIB, based on predominant expression of the corresponding myosin heavy chain genes *Myh7*, *Myh2*, *Myh1*, and *Myh4*, respectively. The morphological, functional and metabolic differences are reflected in distinct transcriptional programs, which likely shape how individual fibers and their associated cell types adapt to training.

Single-cell and single-nucleus transcriptomics could in principle resolve this heterogeneity, but the application to skeletal muscle has been hindered by technical challenges. Standard dissociation protocols for MMC isolation^12^ introduce strong stress-related artifacts that can mask genuine biological signals^13,14^, while the multinucleated nature of fibers leads to their severe underrepresentation in single-cell datasets, making single-nucleus RNA sequencing (snRNA-seq) necessary for adequate coverage. Conversely, rarer yet highly heterogeneous MMC populations are often underrepresented in snRNA-seq, mainly due to limitations in enriching MMC nuclei during isolation. Consequently, a holistic, unbiased, and temporally resolved framework of how myonuclear and MMC populations respond to acute exercise and long-term training has remained elusive.

Here, we present a high-resolution snRNA-seq atlas of mouse skeletal muscle in trained and sedentary states across a longitudinal post-exercise timeline that overcomes these limitations. By utilizing the probe-based FLEX assay on fixed, sorted nuclei, we circumvent dissociation-induced artifacts and achieve unprecedented granularity across all major muscle cell and MMC populations. Our findings reveal that muscle adaptation is not a uniform program but emerges through coordinated, fiber-type-specific transcriptional trajectories, training-modulated regenerative signatures and dynamic multicellular crosstalk.

## RESULTS

### Fixed single-nucleus RNA profiling captures the cellular landscape of muscle adaptation to endurance exercise

To resolve fiber- and cell type-specific transcriptional responses to exercise, we profiled mice before and after six weeks of voluntary wheel running (Fig. 1a). Training significantly improved treadmill performance (Fig. 1b, Extended Data Fig. 1a) and induced a characteristic increase in *M. soleus* mass ^15^, while total lean and fat mass remained stable (Extended Data Fig. 1b–d). To circumvent stress-related transcriptional artifacts common in enzymatic dissociations^13,14^, we applied the probe-based 10x Genomics Chromium Fixed RNA Profiling (FLEX) assay to fixed nuclei isolated from myofibers and MMCs at 0h, 6h, and 12h following exhaustive exercise (Fig. 1a). Myonuclei were enriched via sorting for Pericentriolar Material 1 (PCM1), a marker of the myonuclear envelope^16,17^ (Extended Data Fig. 2a,b). Compared to conventional 3’ snRNA-seq, the FLEX assay provided superior gene detection per UMI with comparable transcript capture sensitivity (Extended Data Fig. 2c–e), indicating comparable transcript capture sensitivity. Notably, stress markers such as *Hspa1a* and *Junb*, markers of stress signatures in cells dissociated from their physiological niche, were absent in our resting samples (Extended Data Fig. 2f). After stringent quality control, we retained high-quality profiles from over 550’000 nuclei, including all known myonuclear and MMC populations (Fig. 1c–e). Unbiased clustering revealed exercise-induced myonuclear subpopulations (called *Ex0h_1–5* or *Ex6h_1–2*) with heterogeneous myosin heavy chain (Myh) isoform expression (Extended Data Fig. 2g), suggesting both shared and bifurcated responses across fiber types. Using digital myonuclear typing based on relative Myh expression (see Methods), we categorized nuclei into established fiber groups with minimal hybrid populations (Extended Data Fig. 2h), as previously reported^9^. Together, our fixed snRNA-seq approach provides unprecedented resolution of post-exercise fiber- and cell type-specific transcriptional dynamics of trained and sedentary muscle tissue in male mice.

**Fig. 1:**
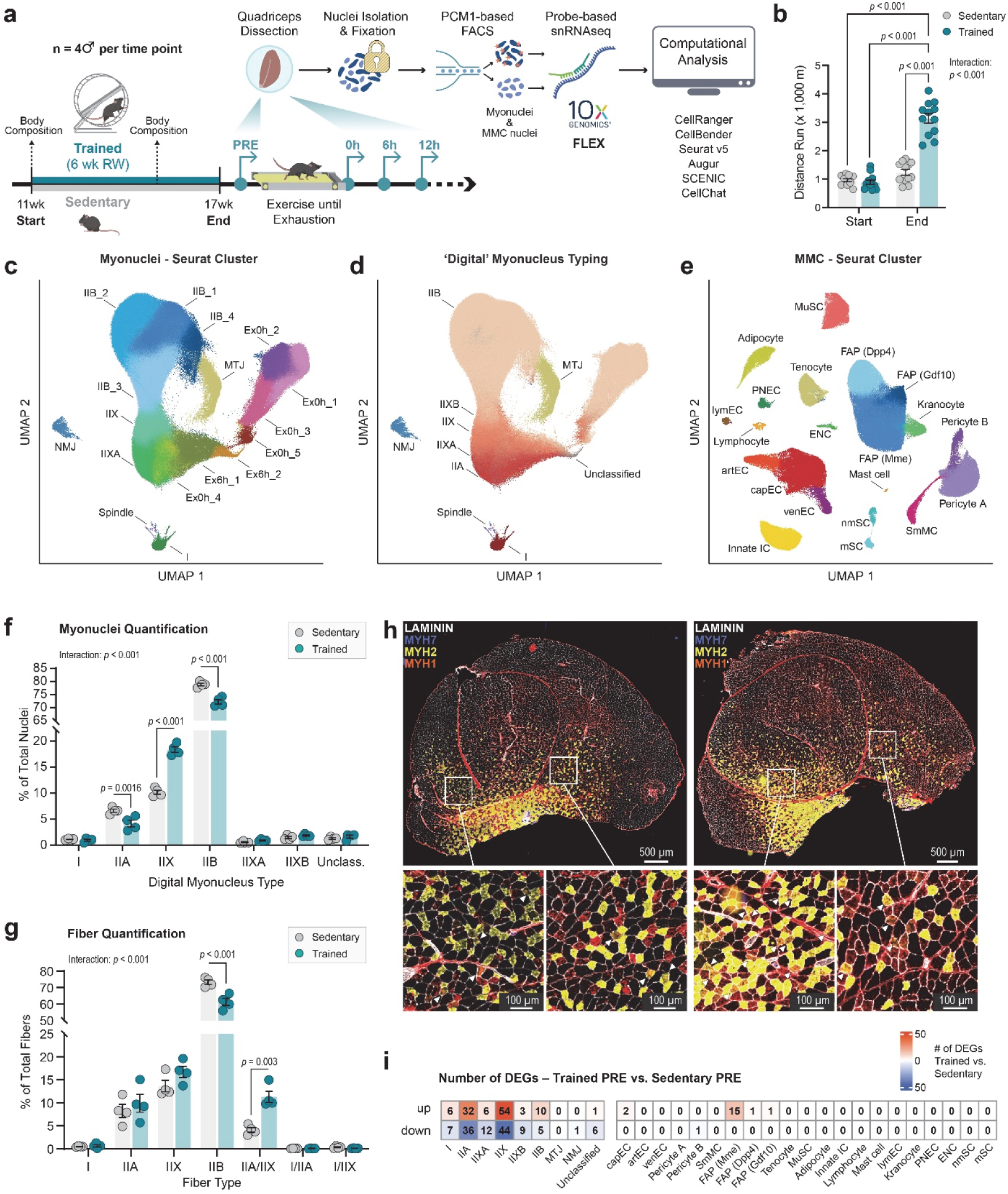
A high-resolution single-nucleus transcriptomic atlas of muscle adaptation to endurance exercise. **a**, Scheme of experimental setup. Control without acute exercise (PRE), mononucleated cell (MMC). Figures are in part generated with BioRender.com. **b**, Distance run during maximal endurance capacity test on the treadmill. Data are mean ±sem, n = 12 (4 per time point excluding control groups to avoid acute exercise effects), Two-way ANOVA with Tukey’s post hoc test with multiple comparisons. **c**, Unifold Manifold Approximation and Projection (UMAP) plot representing the PCM1-positive myonuclear fraction with unsupervised *Seurat* clustering. **d**, UMAP of myonuclear fraction with supervised clustering based on myonucleus typing, guided by the type-specific expression of myosin heavy **e**, UMAP representation of PCM1-negative MMC fraction with unsupervised *Seurat* clustering. **f**, Percentage of total myonuclei of each cluster derived by myonucleus typing (n = 4). **g**, Quantification of immunohistological fiber typing using antibodies against MYH7 (I), MYH2 (IIA), and MYH1 (IIX) showing percentage of total fiber count for each fiber type (n = 4). **h**, Immunofluorescent fiber typing of *M. quadriceps* sections. Blue = MYH7 (I), Yellow = MYH2 (IIA), Red = MYH1 (IIX), Unstained = IIB; arrows indicate hybrid IIA/IIX fibers. Insets show magnified regions highlighting fiber type differences. Scale bars, 500 μm (main) and 100 μm (insets). **i**, Number of significantly up- or downregulated genes per cluster between trained and sedentary muscle. Genes with |log_2_FC| > 1 and *q* < 0.01 were counted as significant. Unless otherwise indicated, data are mean ±sem. Significance for non-expression data was calculated using Two-way ANOVA with Tukey’s post hoc test. Significance for gene expression data was measured from sample- and cluster-wise aggregated pseudobulk counts (n = 4) using *DESeq2* (Wald-test) with *p*-value adjustment using Benjamini-Hochberg.

### IIX myonuclei expand in trained muscle via IIA/IIX hybrid fiber formation

Repetitive bouts of endurance exercise induce well-characterized fiber type transitions (IIB→IIX→IIA) in skeletal muscle^2^ yet how these adaptations manifest at the myonuclear level remains unclear. To address this gap, we quantified myonuclear populations in *M. quadriceps* of trained and sedentary mice at baseline. Remarkably, *Myh1*-expressing (IIX) myonuclei increased ∼1.8-fold in trained muscle (from 10.17% to 18.35%; *p* < 0.001), while *Myh2*-expressing (IIA) and *Myh4*-expressing (IIB) nuclei decreased to ∼0.6 fold (from 6.62% to 4.16%; *p* = 0.0016) and to ∼0.9-fold (from 78.78% to 72.16%; *p* < 0.001), respectively (Fig. 1f). Stable Myh transcript levels across all isoforms (p>0.05) confirmed that these shifts reflect altered myonuclear composition rather than changes in individual nuclear Myh output (Extended Data Fig. 3a). Using classical immunohistological fiber typing of *M. quadriceps* sections (staining for I, IIA and IIB, with unstained fibers assigned as IIX), we confirmed the expected shift toward oxidative types: IIA fibers increased ∼1.8-fold (11.14% to 20.82%, *p* < 0.001), IIX fibers increased ∼1.3-fold (13.27% to 17.57%; *p* = 0.018), and IIB fibers declined to ∼0.8-fold ( 74.75% to 60.71%; *p* < 0.001) (Extended Data Fig. 3b,c). However, because this approach fails to detect IIX hybrid fibers, it likely underestimates IIX fibers. Accordingly, alternative staining for I, IIA and IIX, with unstained fibers assigned as IIB, revealed that this oxidative shift was primarily driven by IIX/IIA hybrid fibers, which increased ∼2.8-fold (4.09% to 11.3%; *p* = 0.003), whereas IIB fibers again decreased to ∼0.8-fold (73.21% to 61.24%; *p* < 0.001) (Fig. 1g,h). Minimal Feret diameter analysis excluded hypertrophy as a driver of IIX myonuclear expansion, as only pure IIA fibers increased in size (interaction *p* < 0.001) (Extended Data Fig. 3d).

These compositional shifts may also promote a more oxidative baseline phenotype, as IIX myonuclei displayed oxidative transcript signatures resembling IIA nuclei, including mitochondrial complex, TCA cycle, and fatty acid oxidation genes, while maintaining intermediate glycolytic, calcium-handling, and lactate metabolism transcripts similar to IIB nuclei (Extended Data Fig. 3e). Differential gene expression (DEG) analysis between trained and sedentary mice of each population at rest, i.e. not affected by an acute exercise bout, revealed few training-induced changes within individual populations, with most occurring in IIA or IIX nuclei (Fig. 1i and Extended Data Fig. 3f). This aligns with the the limited expression changes observed in bulk sedentary vs. trained muscle at rest^2^. Together, these results uncover an unexpected layer of endurance training adaptation, marked by a selective enrichment of transcriptionally oxidative IIX myonuclei in IIA/IIX hybrid fibers.

### Training status dictates fiber- and cell-type-specific transcriptional responses to exercise

Chronic exercise remodels muscle morphology and function^18,19^, yet persistent transcriptional changes remain rare at rest, as seen in both our snRNA-seq data and prior bulk analyses^2^. Nevertheless, sedentary and trained muscles respond differently to an acute exercise bout^2^. To decode such a training state-dependent programming at fiber- and cell type-level, we profiled the transcriptional response to an exhaustive acute exercise bout in trained and sedentary mice. Immediately post-exhaustion (0h), nearly all myonuclei shifted into exercise-specific activation clusters (“*Ex0h_1–5*”, “*Ex6h_1–2*"), whereas MMCs showed only partial recruitment (Fig. 2a and Extended Data Fig. 4a). In sedentary mice, glycolytic IIB myonuclei returned to baseline within 6h, while oxidative IIA and IIX myonuclei remained perturbed even at 12h (Fig. 2b). Conversely, oxidative myonuclei in trained mice recovered significantly faster (IIAX and IIA: interaction p<0.001), mirroring the rapid recovery of glycolytic fibers. This was driven by a reduction in the number of nuclei transitioning into the "Ex6h_2" state at 6h and 12h (interaction p=0.029) (Fig. 2b).

**Fig. 2:**
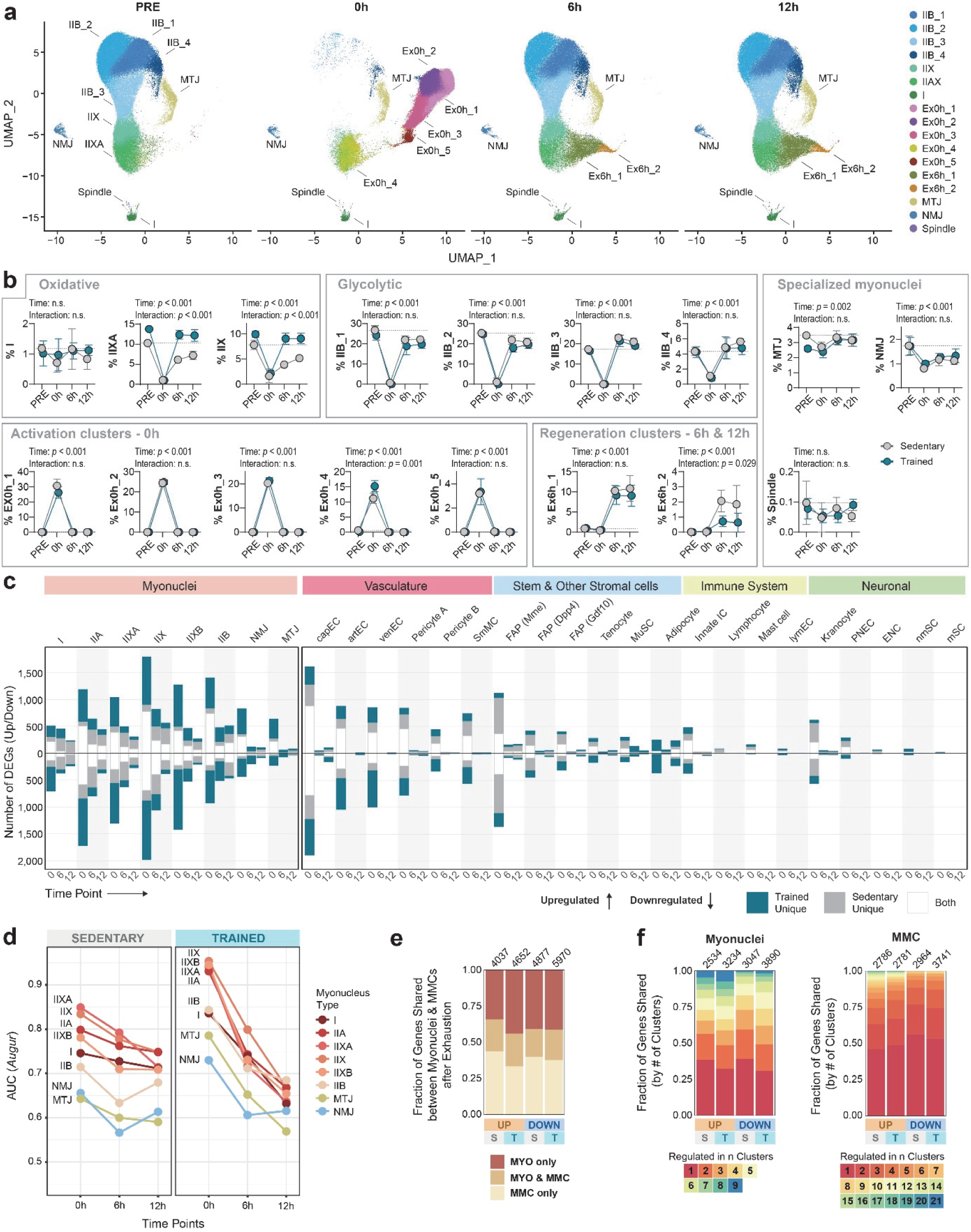
Training remodels the temporal exercise response across muscle cell populations. **a**, UMAP visualization of all myonuclei across four time points (PRE, 0h, 6h, 12h), colored by unsupervised *Seurat* clustering, showing myonuclear subtypes and exercise-induced clusters. **b**, Training state-dependent temporal dynamics of myonuclei subtypes across time points. Data are mean proportion ±sd at each time point. Two-way ANOVA with Tukey’s post hoc test; n = 4 mice per group. Dashed line indicates mean of sedentary baseline (PRE) proportions. **c**, Bar plot summarizing differentially expressed genes (DEGs) between trained and sedentary groups over time, stratified by myonucleus and cell type, and colored by training-state specificity. DEGs were identified via *DESeq2* (Wald-test) on pseudobulk counts aggregated by biological replicate with *p*-value adjustment using Benjamini-Hochberg method. Genes with |log_2_FC| > 1 and *q* < 0.01 were counted as significant. **d**, Line plots showing *Augur*-inferred perturbation scoring (AUC) of myonuclei subtypes across time points and training states. **e**, Stacked bar plots indicating the overlap of DEGs with myonuclear (MYO), mononucleated (MMC), or shared signatures separated by direction of regulation and training-state. DEGs from all time points were merged per training state. S, Sedentary; T, Trained. **f**, Stacked bar plot showing the fraction of up- and downregulated DEGs shared across clusters within the myonuclear (left) or MMC fraction (right). Color intensity reflects cluster-specificity. DEGs from time points were merged. S, Sedentary; T, Trained.

Pseudobulk DEG analysis revealed a robust 0h transcriptional response in both myonuclei and MMCs (|log_2_FC| > 1, *q* < 0.01) (Fig. 2c). While myonuclear DEGs often persisted through 12h, MMCs showed a sharp and sustained decline by 6h, with the exception of adipocytes, which showed increased transcript numbers at 12h. Training induced many novel genes, absent in sedentary muscle, across all myonuclear types, whereas this effect was weaker in MMC and accompanied by further suppression in fibro-adipogenic progenitors (FAPs) (Fig. 2c).

Comparison of DEG overlap across time points revealed that the early 0h response differed markedly from the 6h and 12h response (Extended Data Fig. 4b,c). Cluster-specific perturbation analysis showed that oxidative (particularly IIX) myonuclei were most strongly affected population post-exercise. This perturbation was further amplified by training in all myonuclear types (Fig. 2d). Thus, while training heightens the immediate transcriptional responsiveness to exercise, it simultaneously accelerates homeostatic recovery. In contrast, MMCs exhibited minor training-dependent shifts, characterized primarily by reduced FAP activation and increased responsiveness of resident immune cells and adipocytes in trained muscle (Extended Data Fig. 4d). Comparative DEG analysis revealed a clear compartment-specific logic: only ∼20% of DEGs were shared between myonuclei and MMCs (Fig. 2e). Gene Ontology (GO) analysis supported this dichotomy, linking myonuclear DEGs to regulation of membrane potential and synapse, and MMC DEGs to angiogenesis and inflammation (Extended Data Fig. 4e,f). Shared signatures were limited to general stress responses, such as protein folding and apoptotic signaling, suggesting common stress activation but dissimilar downstream consequences (Extended Data Fig. 4g). Even within each compartment, responses remained highly specialized, with over 50% of myonuclear DEGs restricted to one or two types, and even greater specificity in MMCs (Fig. 2f). Together, these findings demonstrate that training status shapes a highly compartment specific, coordinated response to acute exercise, defined by both heightened responsiveness and enhanced resilience.

### Contextual transcriptional networks drive functional fiber type adaptation

While the preceding analyses demonstrated how training alters transcriptional responsiveness across myonuclear populations, the underlying functional adaptations remained unclear. To define the functional consequences of training-induced plasticity, we clustered DEGs into modules based on their directionality and training-dependency at the 0h post-exercise peak (Fig. 3a–f and Extended Data Fig. 5a–h).

**Fig. 3:**
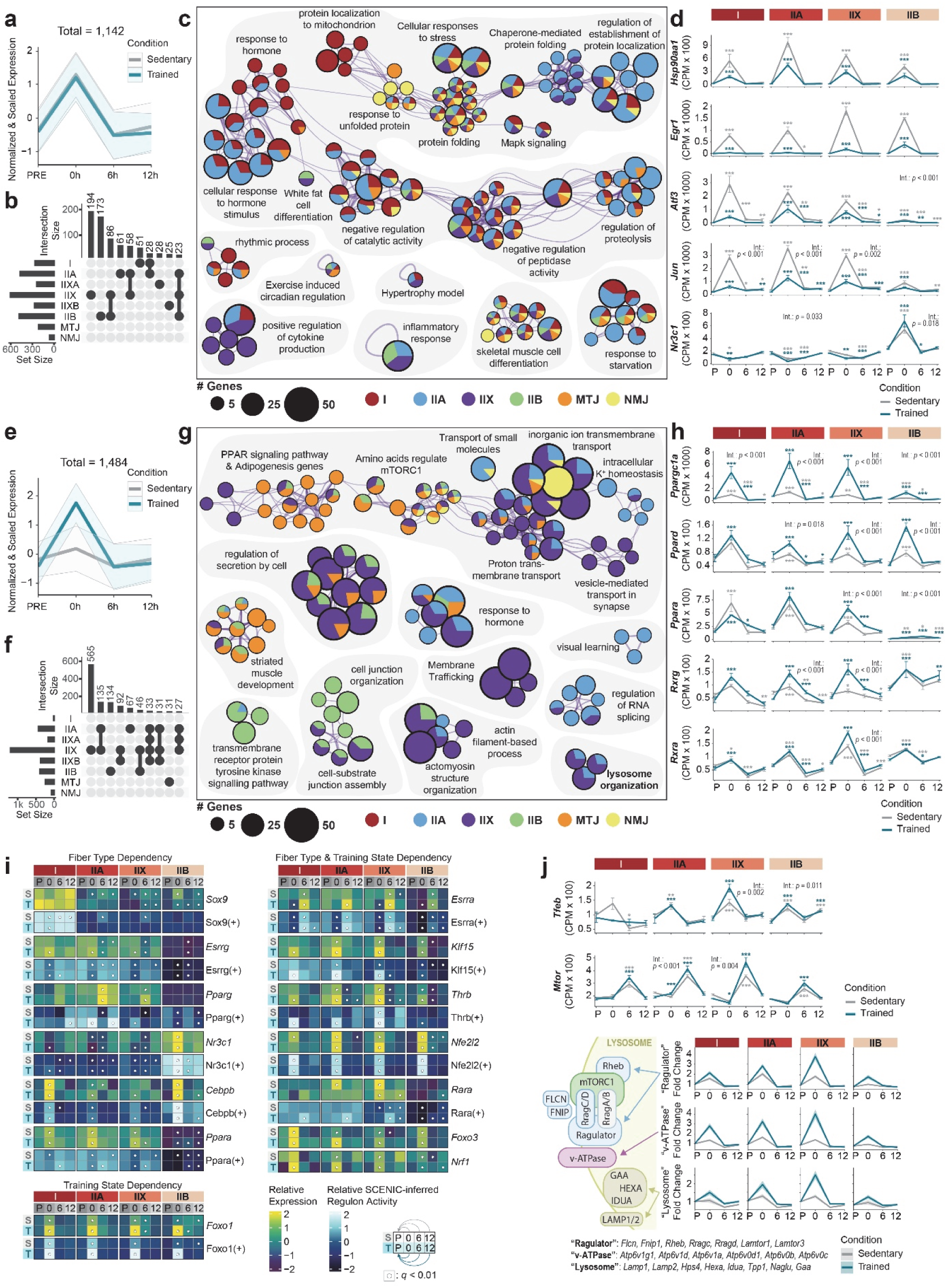
Training reprograms transcriptional networks from general stress response to contextual functional adaptation. **a**, Line plot displaying normalized and scaled expression of a training-state independently activated gene module at the 0h time point. Modules showing similar temporal dynamics across subtypes were grouped. Line plot data represent the mean of module-level averages across subtypes ±sd. **b**, UpSet plot showing overlap of training state-independently upregulated genes at 0h across individual myonuclear clusters. **c**, Enrichment analysis network for training state-independently upregulated genes at 0h. Node size reflects gene set size; circles are colored by contributing myonucleus subtypes. **d**, Line plots showing temporal expression patterns of selected stress-related genes across myonuclei subtypes. **e**,**f**, As in **a**,**b** but for genes with enhanced induction in trained compared to sedentary myonuclei. **g**, Enrichment analysis networks for training-enhanced genes at 0h. Node size reflects gene set size; circles are colored by contributing myonucleus subtypes. **h**, Line plots showing temporal expression patterns of selected training-enhanced transcriptional regulators across myonuclei subtypes. **i**, Heatmaps showing expression levels and SCENIC-inferred transcription factor regulon activity of PGC-1α interaction partners across all time points in major myonucleus subtypes. Dots represent significant changes in comparison to condition-intrinsic PRE controls. S, Sedentary; T, Trained. **j**, Expression trajectories of genes involved in nutrient-sensing via mTORC1 at the lysosome, including *Tfeb* and *Mtor*. Genes were grouped based on functional compartments as indicated. For gene sets, a module depicting the average expression ±sd is shown. For **d**,**h**,**i**: Data are mean ±sem from pseudobulk profiles (n = 4). Statistical significance assessed by *DESeq2* (Wald-test) comparing time point expression profiles to training state-specific control; *p-*values adjusted with the Benjamini–Hochberg method; \**q* < 0.05, \*\**q* < 0.01, \*\*\**q* < 0.001. Overall interaction via likelihood ratio test.

We first identified a training-independent upregulated module (sedentary log_2_FC > 1, *q* < 0.01; interaction |log_2_FC| < 0.5) (Fig. 3a,b). This module included key transcription factors (*Fos*, *Jun*, *Atf3*, *Egr1*), protein chaperones (*Hsp90aa1*, *Hspa1a*, *Dnaja1*, *Dnajb1*), inflammatory signaling (*Nfkb1*, *Clu*), and glucose metabolism regulators (*Irs2*, *Foxo1*, *Csrp3*) (Fig. 3c). Interestingly, some genes were preferentially induced in specific fiber types (e.g., *Jun*, *Atf3* in oxidative types), whereas others were restricted to certain types (e.g., *Nr3c1* in IIB myonuclei) (Fig. 3d). Although trained muscles showed stronger transcriptional perturbation at 0h, these general stress-response genes were significantly dampened compared to sedentary controls (Fig. 3d). This suggests that training shifts myonuclei toward a more targeted adaptive response, reducing reliance on generic stress programs.

To explore these training-enhanced adaptations, we isolated a module of genes with higher activation in trained mice (interaction |log_2_FC| > 0.5, *q* < 0.01) (Fig. 3e). This module was more fiber type-specific and particularly prominent in IIX myonuclei, where 565 of 1’484 genes were uniquely regulated (Fig. 3f). Principal component analysis confirmed a shift toward a more oxidative transcriptional profile post-exercise in all type II myonuclei, with the strongest training effect in IIX myonuclei (Extended Data Fig. 5i). Functional enrichment analysis of training-enhanced genes revealed specific programs, including the PPAR signaling axis and RNA splicing, structural remodeling and cell junction organization, and enhanced membrane trafficking involving vesicle-mediated transport, and secretion (Fig. 3g).

Central regulators of endurance adaptation, specifically *Ppargc1a* (PGC-1α)^2,20,21^ and its interaction partners (*Ppard*, *Ppara*, *Foxo1*, *Foxo3*, *Rxra/b/g*, *Rora/c*, *Esrrg*, *Thrb*) showed dfiber type-dependent regulation, likely driven by distinct physiological demands (Fig. 3h). Accordingly, analysis of expression levels and SCENIC-inferred regulon activities of additional PGC-1α binding partners, revealed contextualized patterns driven by fiber type, training state, or fiber type-specific adaptation (Fig. 3i). Consequently, downstream genes involved in mitochondrial oxidative metabolism, lipid droplet formation, and ROS handling peaked at 6h, even more prominently in trained muscle (Extended Data Fig. 6).

As a second example of fiber-type-specific adaptation, we identified training-enhanced terms for "mTORC1 regulation" and "lysosome organization". These included *Tfeb*, Ragulator-mTORC1 components (*Flcn*, *Fnip1*, *Rragc/d*, *Lamtor1/3*, *Rheb*), and lysosomal proteins (v-ATPase subunits, *Lamp1/2*) (Fig. 3j). These genes showed maximal activation in trained IIX myonuclei, supported by significant interactions for *Tfeb* (q=0.002) and subsequent *Mtor* (q=0.004) expression (Fig. 3j). This signature likely reflects heightened amino acid sensitivity, facilitating a faster transition from a post-exhaustion catabolic state to mTORC1-dependent anabolism. In summary, endurance training refocuses transcriptional networks away from general stress and toward tailored, fiber-type-specific adaptive programs.

### Training rewires multicellular crosstalk during muscle recovery

Emerging evidence identifies MMCs as important mediators of exercise-induced adaptation^22^. Immune, FAPs, and vascular cells contribute to structural remodeling and metabolic reprogramming^23–32^, prompting us to test whether this multicellular communication is transcriptionally encoded and affected by training.

We used *CellChat*^33,34^ to infer receptor-ligand-based cell-cell communication networks from our snRNA-seq dataset. Myonuclei were classified by fiber type, whereas MMCs were grouped into functional categories: Vasculature, FAPs, Immune, and Glial (see Methods). Kranocytes, an NMJ-associated *Mme*^+^ FAPs were confirmed by RNA FISH for *Adamtsl2* and *Shisa3* (Extended Data Fig. 7a–c) and previous studies^35,36^. Total ligand-receptor interactions reached a nadir at 0h post-exercise before increasing at 6h and 12h, a temporal pattern inverse to the peak transcriptional perturbation at 0h. In trained muscle, interaction counts were consistently lower but followed the same trajectories (Fig. 4a). Stratifying these interactions by cellular origin revealed that MMCs dominate the signaling landscape throughout recovery. Notably, the signaling surge at 6h and 12h was driven almost entirely by MMC-MMC communication (Fig. 4b). Signaling strength analysis confirmed this intrinsic propensity of MMCs for engagement, with FAPs and tenocytes acting as the primary signal emitters, while vascular cells and muscle stem cells (MuSCs) served as the primary recipients (Fig. 4c). In contrast, myonuclei showed limited signaling capacity throughout recovery (Extended Data Fig. 7d).

**Fig. 4:**
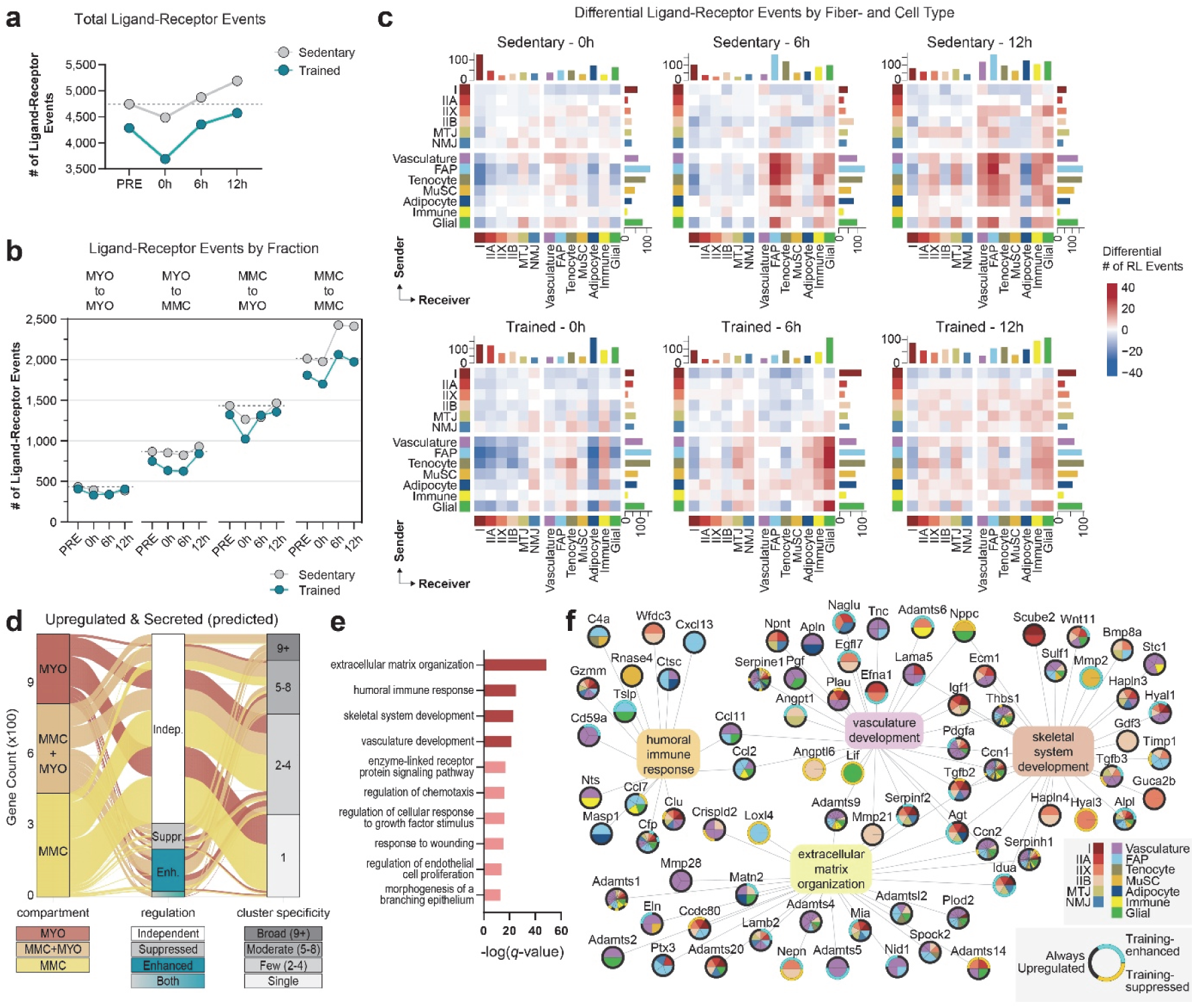
Multicellular communication networks during muscle recovery are mainly driven by MMCs. **a**, Line plot showing the total number of predicted ligand–receptor events across training states and time points, based on *CellChat* inference. **b**, Bar plot quantifying ligand–receptor interactions originating from myonuclei (MYO) or mononucleated cells (MMC), across time points and training states. MMCs are the predominant mediators of multicellular signaling at 6h and 12h post-exercise. **c**, Heatmaps displaying differential ligand–receptor interaction frequencies across cell types in sedentary and trained muscle at 0h, 6h, and 12h post-exercise. Signaling is quantified as differential events compared to respective PRE control. RL: Receptor-Ligand. **d**, Alluvial plot summarizing upregulated secreted factors based on the overlap of DEG analysis (log_2_FC > 0.5, *q* < 0.01) and curated predictions of secreted factors (including Ensembl BioMart, Uniprotkb, GO:Extracellular Space, and the Human Protein Atlas). Genes are shown in relation of cell compartment of origin (MYO, MMC, or both), regulation (training-sensitive or -independent), and cluster specificity (single vs multiple populations). **e**, Gene Ontology enrichment analysis of predicted exercise-induced secreted factors with annotation in at least three of the four reference databases. **f**, Cell type-colored network graph of upregulated secreted genes and their functional associations. Nodes represent secreted factors; edges indicate functional or pathway co-annotation; inner pie chart denotes cell type contribution; outer ring indicates training-dependence. For each GO term, up to 30 genes were selected based on maximal log_2_FC induction (see Methods).

In sedentary mice, the rise in signaling at 6h and 12h post-exercise was widespread across FAPs, vascular, immune, adipocytes, and glial populations, suggesting broad MMC engagement during regeneration (Fig. 4c). In contrast, trained muscles showed fewer differential interactions and a distinct shift toward glial signaling. This likely reflects a state of training-induced re-engagement, as baseline trained muscle displayed reduced communication with glial cells, NMJs, and adipocytes (Extended Data Fig. 7e). Beyond quantitative shifts, exercise and training also altered the qualitative nature of intercellular signaling, as shown by distinct remodeling of major signaling pathways across time and training status (Extended Data Fig. 7f). This is exemplified by the significantly altered myonucleus–to–glia receptor–ligand interactions (log₂FC > 0.5, *q* < 0.01 for ligand or receptor) revealing a dynamic signaling network shaped by time and training (Extended Data Fig. 8a).

A network view of these secreted factors underscores the high degree of cell-type specificity, providing a comprehensive resource for discovering novel myo- and exerkines (Extended Data Fig. 8b). While most factors were induced irrespective of training status, a specific subset showed clear training sensitivity and cell-type restriction (Fig. 4d). GO enrichment linked these factors to hallmarks of structural and functional tissue adaptation, including ECM, immune responses, skeletal system development, and vascular remodeling (Fig. 4e). A network view highlighted the extent and specificity of secreted factor regulation across cell types, offering a resource for discovering novel myo- and exerkines involved in muscle adaptation (Fig. 4f). Together, these findings demonstrate that exercise triggers a delayed, training-dependent multicellular response driven primarily by MMCs, coordinating the remodeling of the muscle microenvironment.

### Training shields oxidative muscle fibers from exacerbated damage

Because multicellular signaling primarily emerged during later recovery, we examined how myonuclear states evolve during the return to homeostasis at 6h and 12h post-exercise, and how these processes are shaped by prior training. Unbiased clustering (Fig. 2a,b) identified two distinct myonuclear populations, *Ex6h_1* and *Ex6h_2*, that emerged exclusively during these later phases. While most myonuclei returned toward baseline after the 0h peak, ∼10% remained in these activated states at 6h, reflecting a spatially bifurcated myonuclear-specific response to mechanical or metabolic burden. Fiber type-specificity of engagement was assessed by quantifying the fraction of IIA, IIX, and IIB myonuclei across time and condition within *Ex6h_1* and *Ex6h_2*, compared to all other myonuclei. In sedentary mice, this transition was heavily biased toward oxidative fibers: ∼53% of IIA and ∼46% of IIX myonuclei entered Ex6h_1, compared to only ∼2% of glycolytic IIB nuclei (Fig. 5a). Entry into *Ex6h_2* was lower overall but again remarkably enriched in oxidative nuclei (Fig. 5a). Training significantly attenuated these transitions of nuclei IIA and IIX myonuclei (IIA-*Ex6h_1*, interaction *p* = 0.009; IIA-*Ex6h_2*, interaction *p* < 0.001; IIX-*Ex6h_1*, interaction *p* < 0.001; IIX-*Ex6h_2*, interaction *p* < 0.001), while IIB myonuclei remained unaffected (Fig. 5a). Compositional analysis showed that >75% of Ex6h_1 nuclei were oxidative, whereas Ex6h_2 contained the highest proportion of unclassified Myh-low nuclei, suggesting that severe perturbation causes oxidative fibers to partially lose their transcriptional identity (Fig. 5b and Extended Data Fig. 9a).

**Fig. 5:**
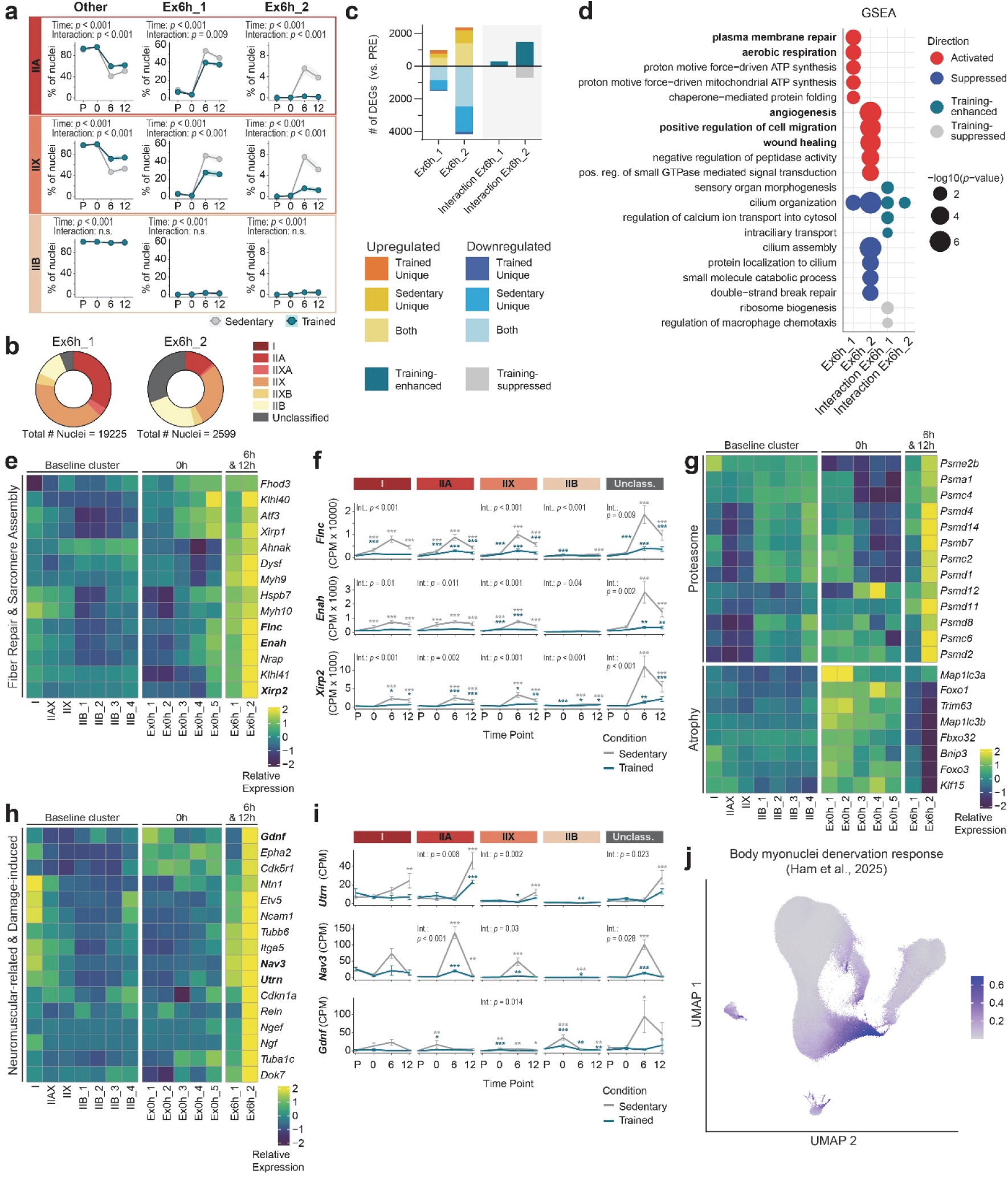
Training remodels bifurcated recovery by restricting entry into highly perturbed regenerative states. **a**, Line plots showing fraction of nuclei in clusters *Ex6h_1*, *Ex6h_2* and all other clusters merged (“*Other*”) across time points and conditions and separated by fiber type. Two-way ANOVA with Tukey’s post hoc test, n = 4 mice per group. **b**, Donut plots showing the myonuclear subtype composition within the *Ex6h_1* and *Ex6h_2* clusters. **c**, Bar plot summarizing the number of DEGs in *Ex6h_1* and *Ex6h_2* compared to PRE baseline, stratified by regulation direction and training-specific interaction effects. Genes with absolute |log_2_FC| > 1 and *q* < 0.01 were counted as significant. Wald-test with *p-*value adjustment based on Benjamini-Hochberg. **d**, Gene set enrichment analysis of DEGs from *Ex6h_1* and *Ex6h_2*, showing functional signatures enriched in trained, sedentary, or both groups. Dot size indicates significance (–log10(*p-*value)); color indicates directionality. **e**, Heatmap showing the expression of selected genes related to sarcomere assembly and repair, including cytoskeletal, scaffold, and membrane repair factors. **f**, Line plots showing temporal expression trajectories of selected regeneration-related genes from (**d**) across myonuclear subtypes. **g**, Heatmap of genes involved in proteasome and muscle atrophy. While proteasome genes are elevated at 6h, canonical atrophy regulators (e.g., *Trim63*, *Fbxo32*, *Klf15*) peak at 0h. **h**, Heatmap of neuromuscular junction (NMJ)-related genes including neurotrophic and structural components elevated in *Ex6h_2*. **i**, Line plots showing temporal expression trajectories of representative NMJ-associated genes. **j**, UMAP with normalized expression of denervation-associated gene set. Top 50 of upregulated genes in body myonuclei post sciatic nerve cut were selected^10^. In **e**,**g**,**h**,**j**: Pseudobulk aggregated counts were normalized for cluster size (CPM) and scaled per row by z-scoring for relative representation. In **f**,**i**: Data represent pseudobulk expression (CPM mean ±sem, n = 4 mice per group). Statistical significance was assessed using *DESeq2* (Wald-test); Benjamini–Hochberg adjusted *p*-values are shown: \**q* < 0.05, \*\**q* < 0.01, \*\*\**q* < 0.001.

To define the functional role of these clusters, we identified 522 and 1’409 training-independent DEGs in Ex6h_1 and Ex6h_2, respectively, compared to baseline (PRE) myonuclei (log_2_FC > 1, *q* < 0.01) (Fig. 5c). *Ex6h_2* displayed additional reprogramming in trained muscle, with 1’474 genes showing positive interaction effects (interaction log_2_FC > 0.5, *q* < 0.01). Gene set enrichment indicated that these clusters drive regenerative and metabolic programs, including plasma membrane repair and aerobic respiration in *Ex6h_1*, and angiogenesis, wound healing and positive regulation of cell migration in *Ex6h_2* (Fig. 5d). In trained muscle, these states were further characterized by "sensory organ morphogenesis" and "cilium organization" signatures, alongside reduced inflammatory markers (Fig. 5d).

*Ex6h_2* was specifically characterized by genes linked to sarcomere assembly and repair^10,37,38^, including cytoskeletal and scaffold proteins (*Enah*, *Flnc*, and *Xirp2*), developmental myosins (*Myh9*, *Myh10)*, and membrane repair factors (*Dysf*, *Ahnak)* (Fig. 5e). An integration of structural and inflammatory cues during regeneration was suggested by the co-expression of inflammatory mediators, including immune receptors (*Csf1r, Edar)* (Extended Data Fig. 9b). These signatures were mainly found in oxidative myonuclei and peaked in unclassified *Myh*-low nuclei (Fig. 5f and Extended Data Fig. 9c), , suggesting that oxidative fibers mount a massive repair response under excessive stress, potentially accompanied by a temporary suppression of *Myh* expression under excessive stress. Crucially, these signatures were significantly attenuated in trained muscle, supporting a protective shielding effect of training against exercise-induced damage (Fig. 5f and Extended Data Fig. 9c).

Although *Ex6h_2* showed evidence of proteolytic remodeling, with strong upregulation of proteasomal subunits (log2FC > 1, *q* < 0.01), canonical atrophy regulators (*Klf15*, *Trim63*, *Fbxo32*) and autophagy genes were not elevated in *Ex6h_1* or *Ex6h_2* but instead peaked immediately after exercise at 0h (Fig. 5g). Trained muscle exhibited an enhanced, transient induction of these atrophy genes at 0h (Fig. 3i and Extended Data Fig. 9d), suggesting that more efficient early-phase "cleanup" may reduce the requirement for prolonged regenerative programs. SCENIC-based inference of transcription factor activity identified elevated activity and partly matching transcript expression of regulators linked to regeneration (*Runx1*, *Smad1*) and inflammation (*Rel*, *Irf5*), alongside less characterized factors such as *Glis3*, *Ets2*, and *Ikzf5* (Extended Data Fig. 9e,f).

Beyond structural remodeling, *Ex6h_2* also upregulated NMJ- and innervation-related genes, including *Utrn*, *Nav3*, and *Gdnf* (Fig. 5h). Some of these NMJ-related genes were dampened in trained muscle, suggesting that training also protects against exercise-induced neuromuscular stress (Fig. 5i). Consistently, a denervation-responsive gene module derived from a sciatic nerve transection dataset^39^ was selectively enriched in *Ex6h_1* and *Ex6h_2* (Fig. 5j). In summary, these data demonstrate that exhaustive exercise forces a subset of oxidative myonuclei into a specialized regenerative state, a fate that is largely circumvented in trained muscle through enhanced resilience and more efficient homeostatic recovery.

### Training protects against exercise-induced NMJ remodeling

Given that *Ex6h_2* myonuclei upregulated NMJ- and innervation-related genes in sedentary muscle, with marked attenuation in trained mice (Fig. 5h–j), we investigated whether these transcriptional changes were accompanied by structural alterations of the NMJ and the state of muscle innervation. We first assessed denervation using neural cell adhesion molecule (NCAM), which is typically restricted to the postsynaptic membrane but becomes cytoplasmic upon impaired innervation. In sedentary mice, the proportion of fibers with mislocalized NCAM increased at 12h and 24h post-exercise before returning to baseline by day 3, whereas trained mice remarkably showed no such changes (Fig. 6a,b). Fibers with mislocalized NCAM were predominantly of type I and IIA (Extended Data Fig. 10a), ), mirroring the enrichment of *Ncam1* transcripts in the two *Ex6h* clusters and the induction in type IIA and IIX myonuclei (Extended Data Fig. 10b,c). Similarly, markers of denervation and NMJ remodeling (*Agrn*, *Musk*, *Myog*, *Lrp4*) were upregulated 24h post-exercise in sedentary muscle while exhibiting a dampened response in trained mice (Fig. 6c).

**Fig. 6:**
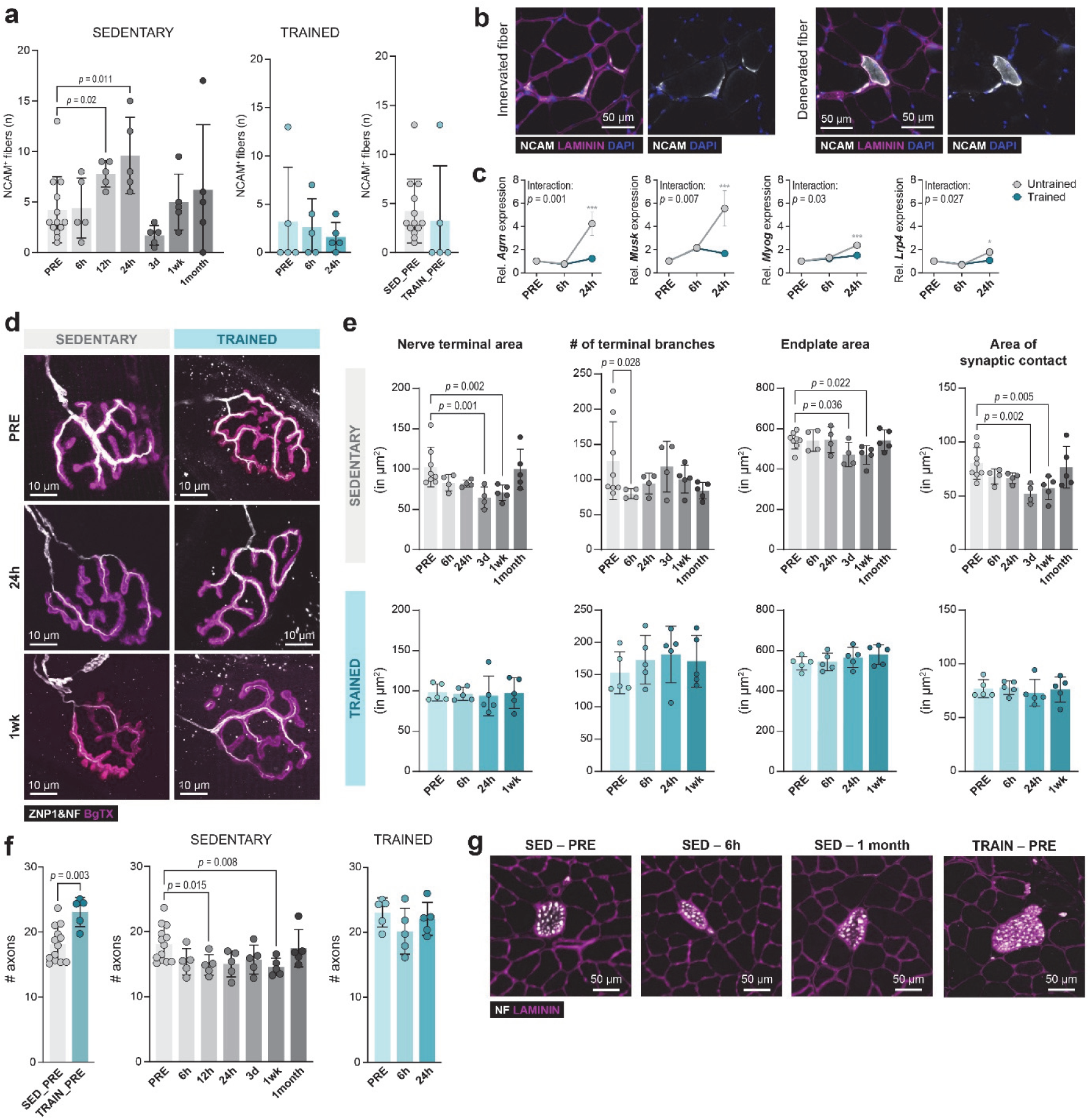
Training preserves neuromuscular junction integrity following acute exercise. **a,** Quantification of NCAM-positive fibers in sedentary and trained mice before and after exhaustive exercise (n = 5 mice/group, except for SED_PRE n = 13 mice). **b,** Representative images of NCAM staining on *M. quadriceps* sections. White = NCAM, Magenta = Laminin, Blue = DAPI. **c**, Quantitative real-time PCR of NMJ-related genes, including *Agrn*, *Musk*, *Myog*, and *Lrp4* of sedentary and trained PRE-, 6 and 24 hours (n = 5 mice/group). **d**, Representative confocal images of PRE and postsynapse of NMJs across groups and time points. Magenta = Bungarotoxin marking Acetylcholine Receptors (AchRs), White = Neurofilament M (NF) and synaptotagmin2 (ZNP1) staining nerve and presynapse. **e**, Quantification of NMJ staining using aNMJmorph analysis measuring nerve terminal area, number of terminal branches, endplate area and area of synaptic contact across groups and time points. Maximum-intensity projections of an average of 23–24 NMJs per mouse were quantified (n = 4–5 mice/group, except for SED_PRE n = 8 mice). **f**, Number of axons in nerve bundles of sedentary and trained mice post-exercise, as well as in baseline comparison (n = 5 mice/group, except for SED_PRE n = 13 mice). **g**, Representative images of neurofilament staining on *M. quadriceps* sections. White = NF, Magenta = Laminin. Data are presented as mean ± SD. One-way ANOVA with Fisher’s LSD post hoc test was used. When data did not pass normality using by the Shapiro-Wilk test, a non-parametric Kruskal–Wallis test with uncorrected Dunn’s post hoc test was applied. An unpaired t-test was used for baseline comparisons. Two-way ANOVA with Tukey’s multiple comparisons test was used for panel **c**. *q < 0.05, **q < 0.01, ***q < 0.001.

Quantitative morphometry of the NMJ (aNMJmorph^40^) revealed that acute exercise in sedentary mice disrupts both pre- and postsynaptic structures (Fig. 6d). Nerve terminal area and perimeter were reduced at 3 days and 1 week post-exercise, whereas terminal branch number declined already at 6h, indicating reduced presynaptic complexity (Fig. 6e and Extended Data Fig. 10d). Concurrently, postsynaptic receptor area and overall endplate area decreased at 3 days and 1 week (Fig. 6e and Extended Data Fig. 10d). These shifts were accompanied by reduced synaptic contact area, pre-postsynaptic overlap and increased NMJ fragmentation, with all parameters returning to baseline within one month (Fig. 6e and Extended Data Fig. 10d,e). Notably, endurance training completely prevented these structural alterations at all time points (Fig. 6e and Extended Data Fig. 10d), despite no differences in NMJ morphology between trained and untrained muscles at baseline (Extended Data Fig. 10e), underscoring a robust protective effect of endurance training. Our snRNA-seq data complement the histological analyses by resolving NMJ-associated transcriptional responses linked to vulnerability in sedentary muscle and protection in trained muscle (Extended Data Fig. 10f).

Since presynaptic alterations preceded postsynaptic remodeling, we examined motor axons after acute exercise. Neurofilament M staining revealed a transient reduction in axon number at 12h and one week post-exercise in sedentary mice, with recovery by 1 month (Fig. 6f,g). In contrast, trained mice exhibited higher axonal counts at baseline compared to sedentary controls, suggestive of increased axonal branching, and maintained this integrity following exercise (Fig. 6f,g). Together, these data demonstrate that exhaustive exercise in sedentary muscle triggers transient presynaptic destabilization and NMJ remodeling. Endurance training confers a robust protective effect, shielding the neuromuscular system from both molecular and structural hallmarks of exercise-induced stress.

## DISCUSSION

Skeletal muscle exhibits remarkable plasticity, underpinning the systemic health benefits of regular exercise^18,19,41–44^. As a syncytium, muscle adaptation emerges from the coordinated activity of spatially and functionally distinct myonuclei and diverse MMCs. While recent efforts have begun to address fiber type-specific adaptations^22,31,45–49^, most have relied on bulk profiling or isolated myofiber analysis. Here, we provide the first snRNA-seq atlas spanning trained and sedentary states across a longitudinal post-exercise timeline. This population-resolved analysis of both myonuclear and MMC compartments reveals previously unrecognized features of the training landscape.

Our data demonstrate that endurance training reshapes the acute transcriptional myonuclear response far more profoundly than the resting transcriptome, consistent with previous bulk analyses^2,50^. While all myonuclear subtypes followed distinct post-exercise trajectories, prior training dampened generic stress programs concomitant with a selective enhancement of fiber-type-intrinsic adaptive responses (Fig. 2c–f and 3a–h). Notably, IIX myonuclei emerged as the most transcriptionally responsive population after an acute bout, and showing the most pronounced training-dependent modulation (Fig. 2d and 3e–g). This heightened responsiveness, coupled with the expansion of IIX nuclei via IIA/IIX hybrid fiber formation (Fig. 1f,g), suggests a model in which training shifts oxidative type II myonuclei towards a particularly adaptive state.

MMCs serve as the primary coordinators of multicellular communication during recovery, exhibiting a highly cell type–specific exercise response (Fig. 2f). Although their early transcriptional perturbation partly mirrored that of myonuclei (Fig. 2c, Extended Data Fig. 4d), their intercellular signaling intensified later, even as most transcriptional activity subsided (Fig. 2c and Fig. 4b,c). This temporal offset suggests a dual role for MMCs: an initial cell-intrinsic response followed by a coordinated phase of tissue remodeling. Previous studies have implicated MMCs in paracrine regulation of fiber function^24,32^, vascular remodeling^23^, immune coordination^27^, and extracellular matrix dynamics^29^. Our dataset provides a foundational framework to dissect the signaling networks that govern paracrine regulation of fiber function, angiogenesis, extracellular matrix and other properties important for shaping muscle recovery after exercise.

A central finding of this study is that post-exercise recovery triggers a bifurcation of a subset of oxidative myonuclei into a late regenerative state. At 6h post-exercise, these myonuclei entered states marked by structural remodeling, proteostasis, inflammation, impaired innervation, and partial loss of fiber identity, suggesting a transient deprogramming process during structural recovery (Fig. 5a,c,e,g,h). Similar transcriptional states have been described in development, aging, and muscular dystrophy^10,37,38^, suggesting that exhaustive exercise engages a conserved program of structural stress. While chronic exercise has been shown to promote damage resilience^2,51^ and support regeneration^52^, our data resolve these dynamics in their specialized myonuclear context and indicate that the burden of recovery falls disproportionately on oxidative myonuclei, with even nuclei of the same syncytium can diverge in transcriptional fate.

This state is closely linked to neuromuscular vulnerability. Direct NMJ analysis in sedentary muscle revealed transient denervation-like response and reversible NMJ remodeling after acute exercise (Fig. 6d–g). Endurance training significantly attenuated both the late regenerative state and these NMJ alterations, indicating enhanced neuromuscular resilience (Fig. 5a and Fig. 6d–g). In line with prior work^53^, the increased abundance of neurofilament-positive axonal profiles at baseline in trained muscle may reflect an enhanced sprouting capacity contributing to protection. More broadly, this NMJ plasticity may not only reflect vulnerability in untrained and resilience in trained muscle, but could also contribute to motor unit rewiring during training adaptation necessary for fiber-type transitions and functional adaptation^19^. Together, these findings suggest that post-exercise perturbation elicits spatially localized regenerative programs within skeletal muscle, including at the neuromuscular junction, and provide a framework to resolve how such microdomain-specific responses shape recovery and adaptation.

Our study has limitations. While we captured the intermediate stages of recovery, the full exercise response likely extends beyond 12 hours^27^. Additionally, the probe-based FLEX assay, while highly sensitive, focuses on protein-coding genes and does not capture non-coding RNA contributions^54–56^. Finally, our data were generated in male mice, *M. quadriceps*, and the context of endurance exercise, and thus do not address muscle-, sex-, or modality-dependent aspects of plasticity^5^.

By providing a population-resolved snRNA-seq atlas of skeletal muscle across training states and recovery, our study demonstrates that endurance adaptation is not a uniform program. Instead, training adaptation emerges from a sophisticated interplay of fiber-type-specific responses, myonuclear fate bifurcation, and multicellular remodeling. This resource offers a map for identifying the molecular pathways that mediate the therapeutic effects of exercise and for understanding the spatial and temporal logic of muscle plasticity.

## Acknowledgments

This study was supported by the Swiss National Science Foundation (Grant no. CRSII5_209252). We thank the FACS Core Facility, the Imaging Core Facility and the Animal Facility of the Biozentrum/University of Basel and the Genomics Core Facility (D-BSSE, ETH Zürich) for their invaluable help with animal caretaking and sample analysis. Data processing and analysis were performed at sciCORE (http://scicore.unibas.ch) scientific computing center at the University of Basel, with support by the SIB (Swiss Institute of Bioinformatics). Some of the graphics were created with Biorender.com.

## Author contributions

C.H., S.D., and M.B. conceptualized and managed the study; S.D. performed snRNA-seq, including sample preparation, library preparation, and snRNA-seq data analysis. P.S. performed data analysis for transcription factor activity prediction; M.B. performed NCAM staining, qPCR, and aNMJmorph experiments. M.B., A.S., S.A.S., G.S. prepared histological samples; M.B., V.A., R.F., S.D. performed and analyzed histological experiments; S.D. performed RNA-FISH; S.D., M.B., V.A., A.S., S.A.S., G.S. performed animal experiments; S.D. and M.B. assembled figures; C.H., S.D. and M.B. wrote the manuscript with the help of all other authors.

### Data availability

Single-nucleus transcriptomics raw data and fully analyzed *Seurat* files have been deposited to GEO under the accession number XXXX. All other data are available upon reasonable request.

## Code availability

The authors declare no original code.

## Materials & Correspondence

Correspondence to C. Handschin

## Declaration of generative AI and AI-assisted technologies in the writing process

During the preparation of the manuscript, the authors used ChatGPT (4o, 5) in order to improve readability. The final content of this manuscript was reviewed and edited as needed and the authors take full responsibility for the content of the publication.

## MATERIAL AND METHODS

### Animals

Male C57BL/6J mice (Janvier Labs) were housed in the animal facility of the Biozentrum, University of Basel, under controlled conditions (22°C, 12-hour light/dark cycle) with ad libitum access to food (standard chow) and water. All experimental procedures were approved by the Cantonal Veterinary Office Basel-Stadt and conducted in accordance with Swiss regulations for animal care and experimentation. Mice were acclimatized to the facility for at least one week prior to handling. At 11 weeks of age, animals were randomly assigned to either a trained or sedentary group. The trained group was provided with voluntary running wheels equipped with hourly distance monitoring over a 6-week period. Sedentary control mice received comparable environmental enrichment in the form of plastic housing structures. Body composition was assessed using an EchoMRI^TM^-100 analyzer in conscious, restrained animals at the start of the running wheel period and after 4 weeks of training.

### Exhaustive exercise treadmill test

A maximal endurance capacity test was performed on a motorized treadmill (5° incline) at the beginning and end of the 6-week training period. Exercise was performed in the morning during the first 4 hours of the light phase to minimize circadian variability. The final test served both as an assessment of endurance adaptation and as an acute exercise intervention, with exhaustion marking the 0 h time point for molecular profiling. Running wheels were removed 48h prior to treadmill acclimatization to minimize acute effects of voluntary activity. Mice were acclimatized to the treadmill over two consecutive days with 5 minutes each at 0, 5, and 8 m/min for the first day and 5 minutes each at 5, 8, and 12 m/min on the second day. On the third day, mice underwent the maximal exercise protocol, starting with 5 minutes at 5 m/min, followed by 5 minutes at 8 m/min, then an increase of 2 m/min every 3 minutes until reaching 32 m/min. Then, speed was increased by 2 m/min every 10 minutes until exhaustion. Mice were considered exhausted when they were unable to resume running despite mild electrical stimuli (0.5 mA, 200 ms pulses at 1 Hz) and gentle prodding. Animals were sacrificed either immediately (0h), 6h or 12h after exhaustion. Muscles were removed, weighted, snap frozen in liquid nitrogen and stored at -80°C until further analysis. Muscles for histology were embedded in OCT and frozen in isopentane at -160°C. Mice assigned to the control “PRE” time point (from both sedentary and trained groups) were not subjected to the treadmill test to avoid confounding acute transcriptional responses.

### Nuclei Isolation and FACS

Matrix-free nuclei isolation from frozen skeletal muscle tissue was established for subsequent 10X Genomics Flex processing. In short, 50 mg of frozen and crushed *M. quadriceps* was lysed using a bead-based tissue homogenizer and lysis buffer containing 10 mM NaCl, 3 mM MgCl2, 10 mM TRIS, Protease Inhibitor (Roche - cOmplete), 0.1 % Triton X-100, 1 µM DTT, and 1 U/µl RNAse inhibitor (Promega - RNasin). Nuclei suspension was filtered through 70 µm and 30 µm cell strainers with a washing and centrifugation step (500g, 5 min, 4 °C) after each filtering using a Wash Buffer with 2 % BSA and RNAse inhibitor (0.2U/µl) in PBS. The final nuclei pellet was fixed for 1h at room temperature and quenched using the 10X Genomics Fixation Kit (1000497). Afterwards, fixed nuclei were stained with Hoechst 33342 and an APC-conjugated anti-PCM1 antibody (#HPA023370, conjugated with Lightning-Link Alexa Fluor-647 Kit #336-0010) for 30 min at 4°C. After final washing, the nuclei suspension was sorted for PCM1-positive and –negative fractions using a BD Aria Fusion Sorter. A small sample of sorted nuclei was taken for imaging. Sorted nuclei fractions where prepared for cryo-storage according to the 10X Genomics Flex protocol and stored at -80°C until the day of sequencing library preparation.

### Single-nucleus library preparation

Single-nucleus RNA-seq libraries were generated using the Chromium Next GEM Fixed RNA kit (10X Genomics, User Guide Rev D CG000527). Nuclei were quantified using Acridine Orange/Propidium Iodide staining and counted with a LUNA-FL fluorescence cell counter. Following probe hybridization, samples were pooled for equal representation (pool and wash option). A total of 16 samples were multiplexed per run, each starting with at least 80’000 nuclei. One biological replicate from each group was included in every run. Myonuclei and muscle-resident mononucleated cell (MMC) nuclei libraries were prepared separately. Sequencing library quality was confirmed via Bioanalyzer. Sequencing was done with a NovaSeq6000 (Illumina) targeting 15’000 reads per nucleus.

### Fiber typing via immunofluorescence

To determine muscle fiber type composition, two immunofluorescence staining protocols were performed in parallel on consecutive 10 μm cryosections of the *M. quadriceps*. One protocol targeted MYH7 (I), MYH2 (IIA), and MYH4 (IIB), with fibers negative for all three classified as IIX. The second protocol targeted MYH7 (I), MYH2 (IIA), and MYH1 (IIX), with triple-negative fibers classified as IIB. Sections were air-dried for 10 minutes at room temperature, followed by 5 minutes of rehydration in PBS. Blocking was performed for 30 minutes at room temperature in PBS containing 0.4% Triton X-100 and 10% goat serum. After three washes in PBS (5 minutes each), sections were incubated for 1h at room temperature with a primary antibody mix diluted in PBS + 10% goat serum. The mix for the MYH4-staining protocol contained BA-D5 (anti-MYH7, DSHB, 1:50), SC-71 (anti-MYH2, DSHB, 1:200), and BF-F3 (anti-MYH4, DSHB, 1:100), whereas the MYH1-staining protocol contained BA-D5 and SC-71 with 6H1 (anti-MYH1, DSHB, 1:25). After primary incubation, sections were washed in PBS (3 × 5 min), followed by a 1h incubation with the appropriate secondary antibody mix in PBS + 10% goat serum. The MYH4 protocol used Alexa Fluor 568 goat anti-mouse IgG1 (A21124, Invitrogen), Alexa Fluor 488 goat anti-mouse IgM (A21042, Invitrogen), and DyLight 405 goat anti-mouse IgG2b (115-475-207, Jackson ImmunoResearch). The MYH1 protocol was identical except that Alexa Fluor 647 goat anti-mouse IgM (A21238, Invitrogen) was used in place of Alexa Fluor 488. All secondary antibodies were used at 1:100 dilutions. After final PBS washes (3 × 5 min), sections were dehydrated in 70% ethanol (2 × 2 min) followed by 100% ethanol (2 × 2 min), mounted with mounting medium, and imaged using a Zeiss Axio Scan.Z1 slide scanner. For fiber type quantification, cryosections were selected at comparable anatomical depth across all samples. Automatic muscle fiber detection was performed using a customized version of the “1_identify_fibers.py” script from the *Myosoft* Fiji toolkit (Hyojung-Choo/Myosoft), adapted for in-house use. Following segmentation, individual fibers were manually classified based on their staining in all fluorescence channels. Morphometric parameters, including cross-sectional area and minimal Feret diameter, were extracted with Fiji.

### RNA-FISH

RNA-FISH was performed on 10 μm cryosections of *M. quadriceps* using the RNAscope Multiplex Fluorescent v2 assay (ACD Bio) according to the manufacturer’s standard protocol. Briefly, sections were fixed with 4% paraformaldehyde for 10 minutes on ice, rinsed with PBS, and dehydrated through graded ethanol. Protease IV treatment was applied for 5 minutes, followed by probe hybridization, signal amplification, and fluorophore development as described in the user manual. Subsequently, immunofluorescence staining was performed to label either the basal lamina or neuronal structures using an anti-Laminin antibody (ab11575, Abcam, 1:200) or an anti-Neurofilament H antibody (AB1991, Merck, 1:200), respectively. Sections were rinsed twice with TBST (TBS containing 1:20,000 Tween-20, pH 7.6), blocked for 1h at room temperature in TBS supplemented with 10% goat serum and 0.1% BSA, and then incubated overnight at 4°C with the primary antibody diluted in TBS + 0.1% BSA. After three washes with TBST, sections were incubated for 30 minutes at room temperature with the secondary antibody (Alexa Fluor 750 goat anti-rabbit IgG, A21039, Invitrogen; 1:500 dilution). Following final TBST washes, sections were mounted with DAPI-containing mounting medium and imaged.

### NMJ-staining and quantification

GC muscles were dissected and stretch-pinned in a petri dish containing 4% paraformaldehyde (PFA) in PBS for 30 minutes at RT. After PBS washes, muscles were placed under a stereomicroscope and gently separated into bundles of approximately 10–15 fibers by pulling from the tendons with fine tweezers. Bundles were incubated in 100 mM glycine for 1h, permeabilized in 0.5% Triton X-100 (4 x 15min), and blocked for 1h in 4% BSA, 5% goat serum, and 0.5% Triton X-100 in PBS. All the steps were performed in an orbital shaker. Primary antibodies were applied overnight at 4°C in blocking solution: anti-Neurofilament M (pan-axonal; 837904, BioLegend) at 1:200 and anti-Synaptotagmin2 (DSHB; ZNP-1) at 1:200.

The following day, bundles were washed (3 x 15min) and incubated for 2h in the dark with secondary antibodies (AlexaFluor 568 anti-mouse IgG1, A21124, Invitrogen, and AlexaFluor 568 anti-mouse IgG2a, A21134, Invitrogen) at 1:500, together with AlexaFluor 488-conjugated bungarotoxin (B13422, Invitrogen) at 1:1000, diluted in blocking solution. Bundles were washed (4 x 15min) and mounted in ProLong Gold Antifade Reagent (Invitrogen), flattened under load, and allowed to dry before imaging.

Confocal images were acquired on a SpinD microscope (Olympus) using a 100x objective. Maximum-intensity projections of an average of 23–24 NMJs per mouse (n = 4–5 mice/group, except for SED_PRE n = 8 mice) were quantified using aNMJ-morph^1^. Average values were used for statistical analyses.

### NCAM and Neurofilament staining

10 µm frozen quadriceps cross-sections were fixed with 4% PFA in PBS, for 15min RT. After PBS washes (3 x 5min), sections were incubated for 1h at RT with blocking buffer containing 0.5% Triton X-100 and 5% Goat Serum in PBS. Primary antibodies were applied overnight at 4°C in 5% goat serum in PBS: anti-NCAM (5032, Millipore) at 1:100, anti-Neurofilament M (pan-axonal; 837904, BioLegend) at 1:200, and anti-Laminin (ab11576, Abcam). The following day, sections were washed in PBS (3 x 5min) and incubated with for 2h at RT with secondary antibody diluted in blocking solution: AlexaFluor 647 anti-rabbit IgG (711-605-152, Jackson, 1:200), AlexaFluor 568 anti-mouse IgG2a (A21134, Invitrogen, 1:500), and AlexaFluor 488 anti-rat IgG (A11006, Invitrogen, 1:200). After additional PBS washes (3 x 5min), coverslips were mounted using ProLong Gold Antifade Reagent with DAPI (Invitrogen). Images were acquired on a AxioScan (Zeiss) using 20x objective. For NCAM, the total number of NCAM-positive muscle fibers within each cross-section was manually quantified. For neurofilament, every axon bundle in the entire cross-section was examined and the number of individual neurofilament-positive axons contained within each bundle was counted. In this case, for each muscle, two sections located approx. 100 µm apart were analyzed and the average values were used for statistical analyses. For both quantifications, n = 5 mice/group, except for SED_PRE n = 13 mice.

### RNA extraction and qPCR

RNA was extracted using a hybrid method, in which 25mg of frozen crushed quadriceps muscles were homogenized using bead beating (10s at 6.5m/s speed, 100s break, 3 cycles) in 1 ml TRI reagent (Sigma-Aldrich). Lysed tissue underwent phase separation using 1-bromo-3-chloropropane, and the RNA in the aqueous phase was pulled-down with isopropanol and washed with ethanol. RNA samples were further purified using 3M of Na-Acetate and Isopropanol and washed again with 75% ethanol. RNA concentration was measured using Nanodrop-1000 (Thermo Fisher Scientific) and 1 µg of RNA was used for cDNA synthesis using Reverse Transcriptase High-Capacity cDNA Reverse Transcription Kit (Applied Biosystems). qPCR was performed using Fast SYBR Green Master Mix (Applied Biosystems) and QuantStudio 5 (Applied Biosystems). Reactions were done in triplicates with the addition of negative controls. Relative mRNA expression was determined using comparative ΔΔCT method, normalizing to a suitable housekeeper (TATA-box binding protein, TBP). Primers were tested prior usage.

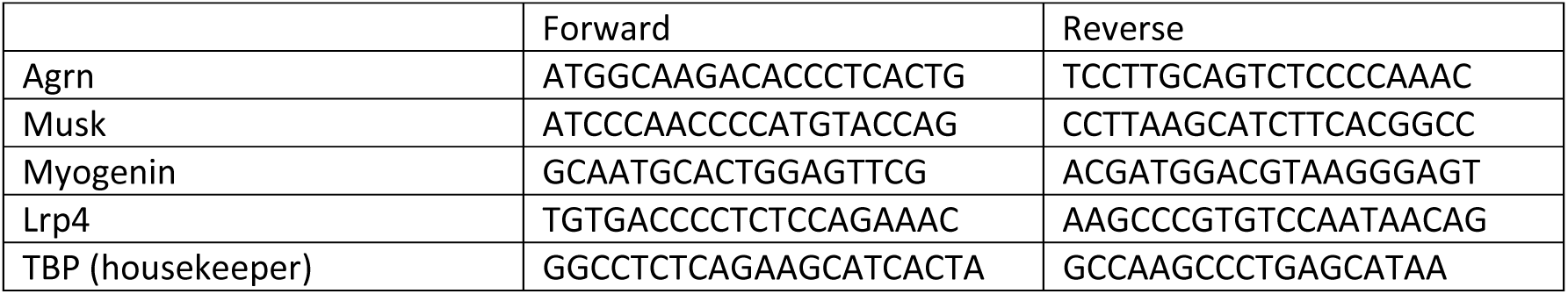

### Data Analysis

#### Preprocessing, quality control, data integration, and clustering

Feature-barcode matrices for each sample were generated using the *multi* pipeline from *CellRanger* (version 7.1.0) using the mouse GRCm39 (mm10) reference genome. Ambient RNA contamination was removed using the *remove-background* function of *CellBender*^2^ (version 0.3.0). Processed count matrices were imported into R (version 4.3.2) for downstream analysis with *Seurat*^3^ (version 5). Initial quality control included filtering out nuclei with fewer than 500 or more than 5’000 detected features, total UMI counts exceeding 15’000, or mitochondrial transcript content greater than 3%. Putative doublets were identified and scored using the *DoubletFinder* algorithm, and samples were subsequently merged, treating myonuclei and MMC nuclei separately. Integration across samples was performed using *Seurat*’s *IntegrateLayers* function with batch correction based on *Harmony*. Clusters enriched for high doublet scores were excluded, followed by reclustering of the cleaned dataset. In the MMC dataset, additional filtering was applied to remove contaminating myonuclei by excluding nuclei with Titin (*Ttn*) expression above a threshold of 1.

#### “Digital” myonucleus typing

To enable fiber type–specific differential expression analysis, myonuclei were classified into subtypes based on the expression of the four major myosin heavy chain (*Myh*) isoforms: *Myh1* (IIX), *Myh2* (IIA), *Myh4* (IIB), and *Myh7* (I). For each nucleus, the total *Myh* expression was calculated as the sum of log-normalized expression values obtained from *Seurat*’s *NormalizeData* function. A myonucleus was assigned to a specific fiber type if at least 45% of its total *Myh* signal originated from a single isoform and the expression level of that isoform exceeded 1.5. Hybrid fiber types were defined when two isoforms each contributed >45% of the total *Myh* expression and both had expression levels above 1.5. Nuclei annotated as specialized subtypes, including neuromuscular junction (NMJ), myotendinous junction (MTJ), and spindle fiber nuclei, were retained under their original annotation. Myonuclei that did not meet any of the above criteria were labeled as “Unclassified”.

### Pseudobulk differential gene expression analysis

Differential gene expression analysis was performed using *DESeq2*^4^ in a pseudobulk framework. For each cell type or myonuclear subtype, raw UMI counts were aggregated by biological replicate (n = 4), condition, time point, and cluster identity. The resulting pseudobulk count matrices were used to construct *DESeq2* datasets, including the interaction term (*∼ condition + time + condition:time*). Prior to model fitting, low-expressed genes were filtered out, keeping genes with cluster-size adjusted minimal counts in at least 4 pseudobulk samples. After filtering, *DESeq2* was run using default settings, followed by log_2_FC shrinkage with the *lfcShrink* function using the “*ashr*”^5^ method to improve effect size estimation. Result tables were extracted for each time point in comparison to the condition-specific “PRE” baseline, as well as the respective interaction terms. To identify genes enriched in the *Ex6h_1* and *Ex6h_2* clusters, we adjusted the existing unsupervised *Seurat* clustering by combining all myonuclei of the “PRE” time point as control cluster (clustering termed “PRE.combined”). We then applied pseudobulk aggregation based on replicates, condition, and the adjusted “PRE.combined” clustering and ran *DESeq2* with the interaction term *(∼ condition + PRE.combined + condition:PRE.combined*), allowing assessment of the main effect in the *Ex6h_1* and *Ex6h_2* clusters, as well as the training-state-dependent differences. Genes with at least 20 counts in 4 samples were kept and log_2_FC was shrunk as before.

Genes were considered significantly regulated in comparisons if they had a baseMean ≥ 10, absolute log_2_FC ≥ 1, and *q* < 0.01. For interaction terms, a log_2_FC threshold of 0.5 was used, with the same criteria for baseMean and *q*-value.

Gene expression heatmaps were generated based on aggregated expression values from *Seurat*’s *AggregateExpression* function, grouped by time point and myonuclear subtype. Replicates were pooled for better heatmap visualization. Differences in sequencing depth and cluster size were normalized by conversion to counts per million (CPM). CPM values were then log-transformed and z-score normalized by gene (row-wise) to highlight relative expression pattern across groups. Heatmaps were generated using *pheatmap*. Line plots were generated by aggregating raw counts per replicate, cluster, and group, followed by conversion to CPM and visualization using *ggplot2*. Mean expression values were displayed across time points for each condition, with error bars indicating the standard error of the mean.

### Definition of gene set modules

Gene modules with coherent expression patterns across time and training were defined by selecting genes according to the following criteria across all myonuclear subtypes. We defined two categories: training-independent genes, which were significantly regulated in sedentary mice without a significant condition interaction, and training-dependent genes, which included sedentary-regulated genes with a significant interaction term as well as all other genes with significant interaction effects (|log_2_FC| ≥ 0.5, *q* < 0.01). For each cluster, aggregated expression matrices were generated and filtered to retain either the training-independent or training-dependent gene sets, resulting in two matrices per cluster.

For each matrix, CPM values were log-transformed, mean-centered, and row-wise z-score normalized. Heatmaps were generated and clustered using k-means to identify distinct temporal expression profiles. Genes belonging to similar expression modules across clusters were then grouped into broader training-independent or training-dependent modules. For each module, average expression across clusters was calculated, and line plots were generated displaying the mean trajectory with ribbons representing the standard deviation across contributing clusters. The overlap of gene identities across myonuclear subtypes within each module was visualized using *UpSet* plots, highlighting the subtype-specific contributions to each expression program. Only nuclei assigned to clearly defined fiber types (I, IIA, IIX, IIB), NMJ, or MTJ subtypes were included; hybrid and unclassified myonuclei were excluded from this analysis.

### Enrichment analysis

Ontology-based enrichment analysis was performed in a myonuclear subtype-specific manner using the *Metascape*^6^ web platform (metascape.org) with default parameters. To account for differences in gene expression repertoires between myonuclear subtypes and MMC types, we provided custom background gene lists whenever applicable. *Metascape* clusters enriched terms by computing pairwise similarity based on shared gene membership and applies hierarchical clustering to group related terms. For visualization, only the representative term of each cluster was displayed. Enrichment network plots generated by *Metascape* were further processed and visually adapted using *Cytoscape (version 3.10.2)*.

To functionally characterize the *Ex6h_1* and *Ex6h_2* clusters, we performed rank-based gene set enrichment analysis (GSEA) using the *gseGO* function from the *clusterProfiler*^7^ R package (version 4.10.1) with the *org.Mm.eg.db* annotation database. For each cluster, GSEA was applied separately to both the main effect (Ex6h vs. PRE myonuclei) and the condition interaction results from the *DESeq2* analysis. Gene lists were ranked by log_2_FC and enrichment was performed against the Gene Ontology (GO) Biological Process ontology with a gene set size between 10 and 500 and a *p*-value cutoff of 0.05. For visualization, we displayed only the top five enriched terms per analysis direction (up- or downregulated) and category (main effect or interaction term).

### Perturbation scoring

Perturbation analysis was performed using the *Augur*^8^ R package (*github/neurorestore/Augur*, version 1.0.3) to quantify transcriptional activation between baseline (PRE) and post-exercise myonuclei and MMC at each time point (0h, 6h, 12h), separately for sedentary and trained mice. *Augur’s calculate_auc()* function was applied to compute area under the curve (AUC) scores per cell type, with permutation-based controls (*augur_mode = ’permute’*) used to establish null distributions. To assess training-dependent shifts in perturbation magnitude, we applied *Augur*’s *calculate_differential_prioritization()* function to compare AUCs between sedentary and trained groups, using 100 subsamples and 1000 permutations. Unclassified and spindle nuclei were excluded from this analysis. AUC trajectories and subtype rankings were visualized across time points for each condition using line plots.

### Transcription factor regulon activity analysis

Transcription factor (TF) activity was inferred using the *pySCENIC*^9^ pipeline (version 0.12.1) with default settings. Raw UMI count matrices were exported from the *Seurat* object and used as input for gene regulatory network (GRN) inference with *GRNBoost2* from the *arboreto* package. TF-target modules were identified using *modules_from_adjacencies*, followed by motif enrichment pruning using *cisTarget* databases (mm10 v10nr) to refine regulons. Regulon activity scores were then calculated using *AUCell*, which estimates the enrichment of each regulon’s targets within the expression profile of individual nuclei. The resulting AUC matrix was imported into *Seurat* as a new assay (“TF.AUC”) using *CreateAssayObject*, enabling visualization of regulon activity across UMAP embeddings and metadata variables via standard *Seurat* functions.

### Cell-cell communication and secreted factor analysis

Intercellular communication analysis was performed using the *CellChat* R package (version 2.1.2), following the standard comparative workflow. Prior to *CellChat* analysis, we merged the processed *Seurat* objects of myonuclei and MMC populations and re-ran the standard *Seurat* preprocessing pipeline, including normalization, integration, and dimensionality reduction. However, we retained the original clustering obtained from the individual datasets to preserve subtype resolution. For improved interpretability in the context of cell–cell communication, MMC clusters were grouped into broader functional categories: Vasculature (capillary, venous, arterial, and lymphatic ECs, pericytes, smooth muscle), "FAP" (*Dpp4*⁺, *Gdf10*⁺, *Mme*⁺), "Immune" (lymphocytes, mast cells, innate immune cells), and "Glial" (myelinating and terminal Schwann cells, perineurial epithelium, endoneurial cells, and neuromuscular junction (NMJ)-capping kranocytes). *CellChat* was run separately for each group defined by condition and time point to enable comparative analysis of signaling networks. The standard *CellChat* pipeline was applied, including inference of overexpressed ligand–receptor pairs, pathway-level communication probabilities, and visualization of dynamic signaling patterns.

To extend beyond curated ligand–receptor pairs, we annotated secreted factors using four databases with secreted factor prediction: Ensembl BioMart, UniProtKB, GO:Extracellular Space (GO:0005615), and the Human Protein Atlas. Proteins with predicted transmembrane domains were excluded from these references whenever possible. We intersected this reference set with all DEGs showing a log_2_FC > 0.5 and *q* < 0.01 across all clusters and time points and displayed in an alluvial plot the cellular origin (myonuclei or MMCs), training-dependence, and cluster specificity of these predicted secreted DEGs. Among these, factors annotated by at least three databases were subjected GO enrichment using *Metascape*. From the top four GO terms, we extracted up to ten most strongly training-enhanced, training-suppressed, and training-independently upregulated genes and visualized them in a cluster-annotated network via *Cytoscape*.

### Statistics

Differential gene expression analyses and time-point-specific interaction effects were calculated using *DESeq2* with the Wald test and Benjamini–Hochberg correction for multiple testing. Overall interaction terms were evaluated using *DESeq2*’s built-in likelihood ratio test (LRT) framework to assess the effect of training condition on the temporal response. Normality and lognormality were assessed for each dataset using the Shapiro–Wilk test. Parametric tests were applied to normally distributed data, and non-parametric tests were used for data that were not normally distributed. Unpaired two-tailed t-tests (parametric Student’s t-test or non-parametric Mann-Whitney) were used for pairwise comparisons of quantitative measurements. Multiple t-tests were applied when assessing multiple pairwise comparisons between two groups. For multi-factorial comparisons (e.g., body composition, myonuclear cluster proportions, endurance performance, and regulon activity), two-way ANOVA followed by Tukey’s post hoc test was used, whereas one-way ANOVA with Fisher’s LSD or Dunn’s post hoc tests was used for comparisons of more than two groups with a single independent variable. Unless otherwise stated, n = 4 mice per group.

---

**Extended Data Fig. 1:**
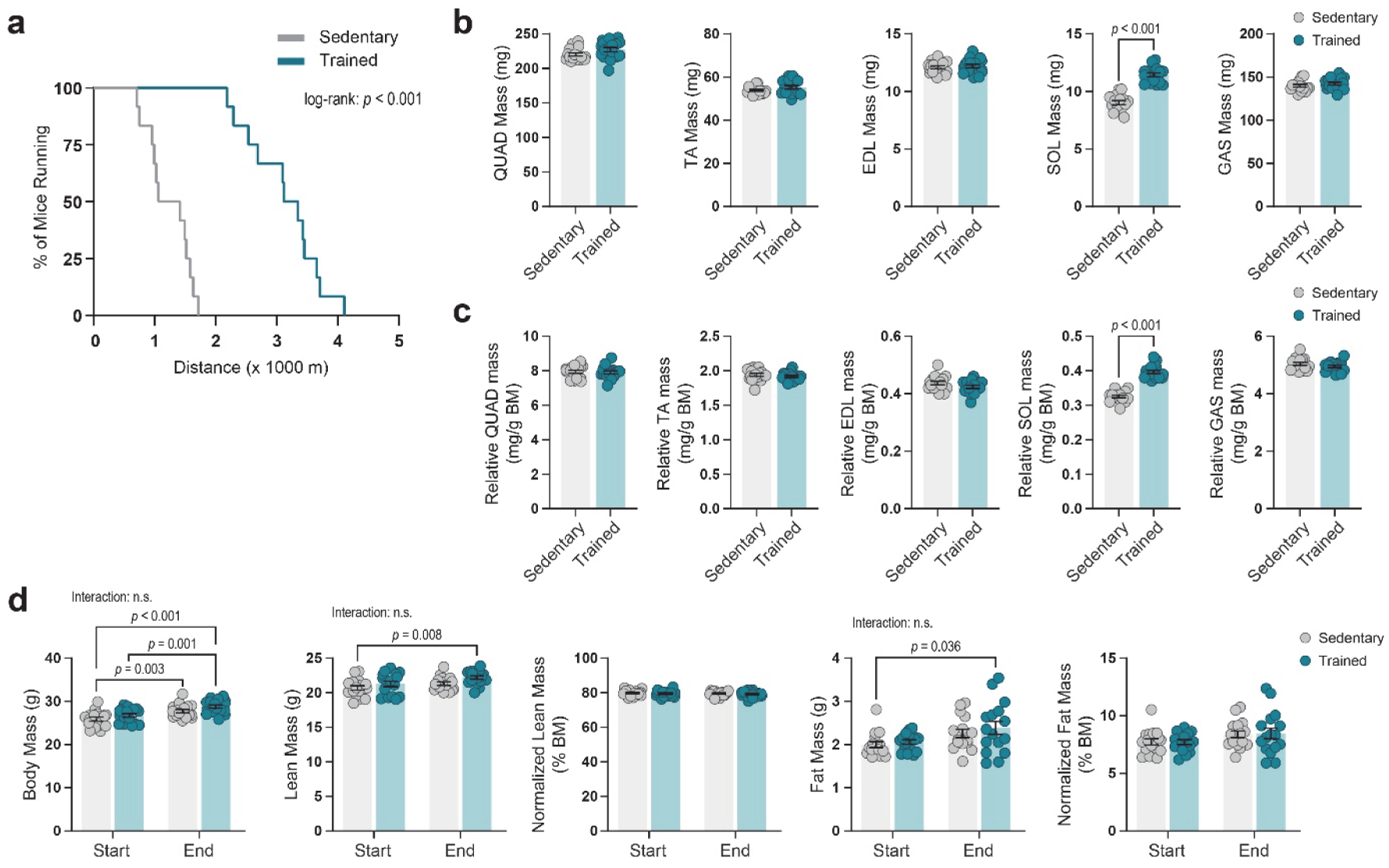
Supplementary cohort data. **a**, Kaplan-Meier survival curves of mice lasting on the treadmill during the final maximal endurance capacity test. Significance calculated by two-sided log-rank test. **b**,**c**, Absolute muscle mass (**b**) and relative muscle mass normalized to body mass (BM) (**c**) of *M. quadriceps* (QUAD), *M. soleus* (SOL), *M. tibialis anterior* (TA), *M. extensor digitorum longus* (EDL), and *M. gastrocnemius* (GAS). Whenever available, average of contralateral muscles was taken. n = 16; data are mean ±sem; unpaired student’s t-test; parametric, when normality confirmed via Shapiro-Wilk, otherwise nonparametric Mann-Whitney test. **d**, From left to right: total body mass, EchoMRI-derived total lean mass, lean mass normalized to total body mass, EchoMRI-derived total fat mass, and fat mass normalized to total body mass. Data are mean ±sem; n = 16 per group, Two-way ANOVA with Tukey’s post hoc test with multiple comparisons.

**Extended Data Fig. 2:**
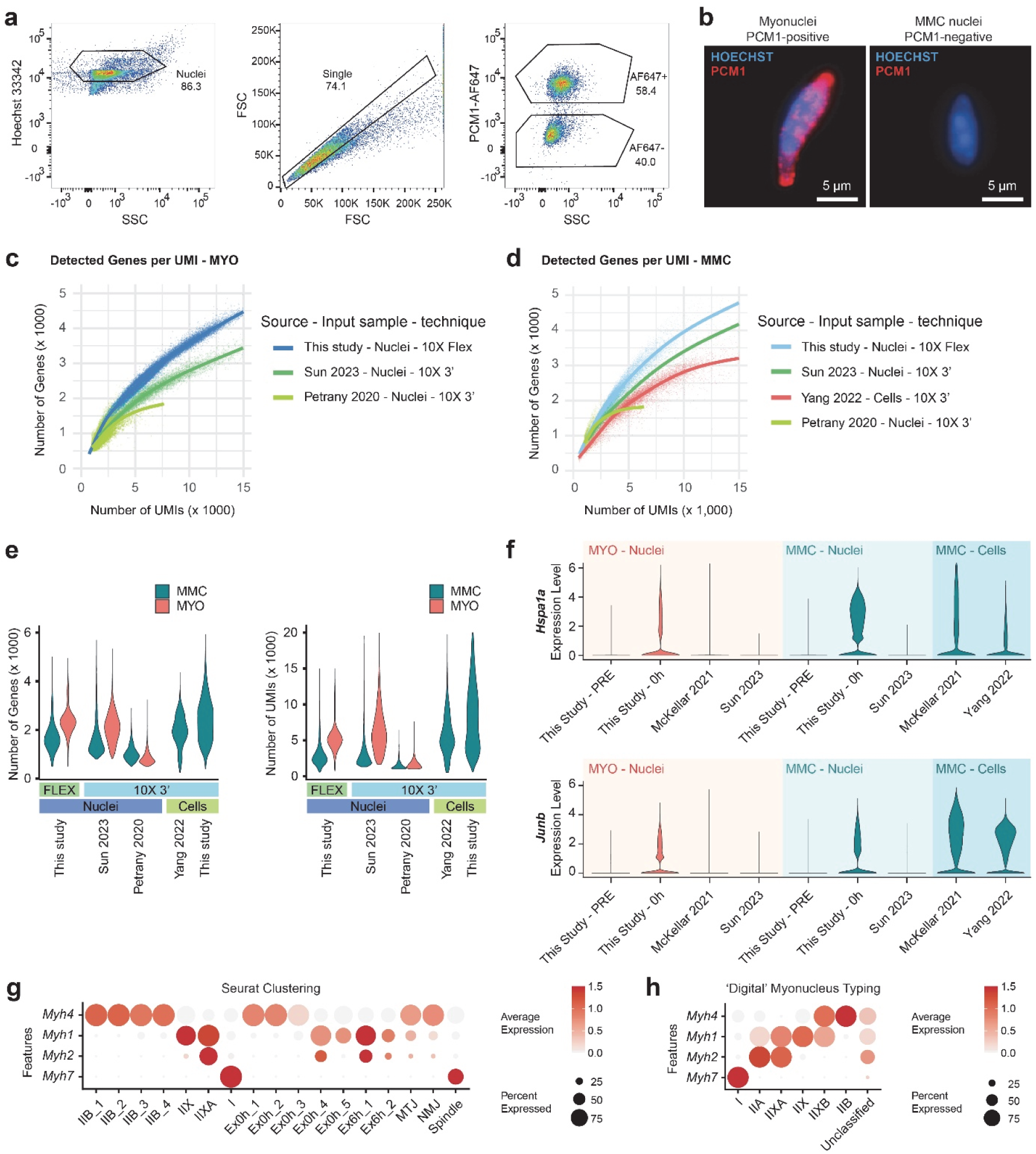
FACS-based nucleus isolation, method comparisons, and digital fiber typing. **a**, Representative scatter plots with gating strategy for PCM1-positive and negative nuclei. Nuclei were sorted based on Hoechst 33342 and APC-conjugated anti-PCM1 antibody. **b**, Immunofluorescent images of FACS-sorted PCM1-positive myonucleus (left) and PCM1-negative MMC nucleus (right) with Höechst 33342-stained DNA (blue) and anti-PCM1 (red). **c**,**d**, Number of genes detected per UMI in myonuclei (**c**) and MMCs (**d**). **e**, Number of detected genes (left) and UMIs (right) per nucleus or cell, comparing data of multiple snRNA-seq and scRNA-seq datasets. **f**, Violin plots with expression levels of stress-related genes (*Hspa1a* and *Junb*) in distinct techniques and sample input types. **g**, Dot plot showing average expression and percentage of myonuclei expressing *Myh* types across unbiased *Seurat* clustering. **h**, Dotplot showing average expression and percentage of myonuclei expressing Myh isoforms across clustering from myonucleus typing.

**Extended Data Fig. 3:**
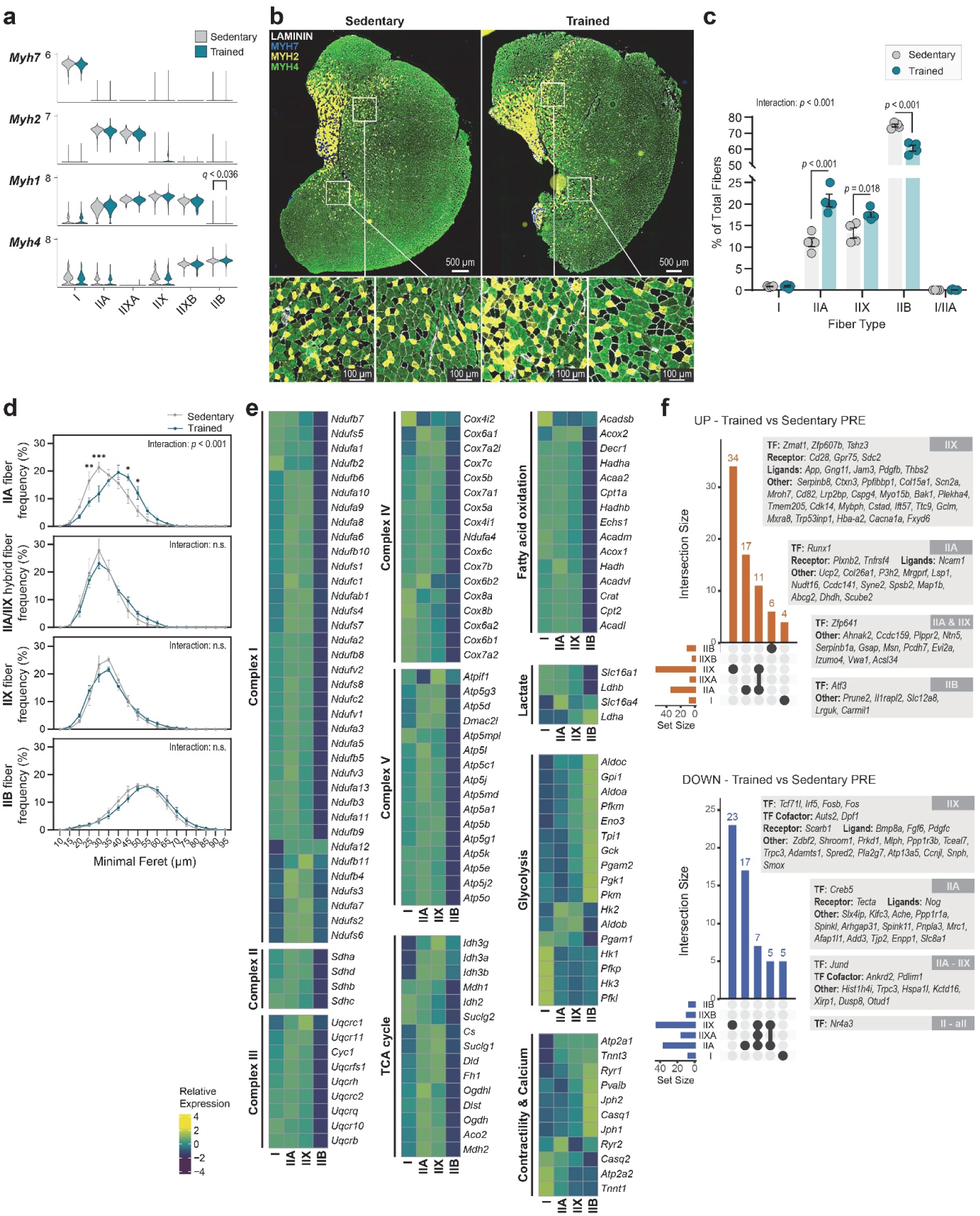
Transcriptional and compositional baseline difference between trained and sedentary muscle. **a**, Violin plots showing *Myh* isoform expression in sedentary and trained muscle by myonucleus type. **b**, Immunofluorescent fiber typing of *M. quadriceps* muscle sections. Blue = MYH7 (I), Yellow = MYH2 (IIA), Unstained = IIX, Green = MYH4 (IIB). Insets show magnified regions highlighting fiber type differences. Scale bars, 500 μm (main images) and 100 μm (insets). **c**, Quantification of immunohistological fiber typing using antibodies against MYH7 (I), MYH2 (IIA), and MYH4 (IIB) showing percentage of total fiber count for each fiber type.. Data are mean ±sem; n = 4; Two-way ANOVA with Tukey’s post hoc test. **d**, Histomorphometry of fiber minimal Feret across fiber types. **e**, Heatmap of relative gene expression (pseudobulk) for mitochondrial and metabolic genes across major fiber types. Expression values counts per million (CPM) for cluster-normalization and are scaled by row. **f**, UpSet plots showing overlap of differentially expressed genes across fiber types (I, IIA, IIX, IIB) at baseline (PRE) in trained versus sedentary muscle. Genes upregulated (left) and downregulated (right) in trained mice are shown with selected transcription factors (TFs), ligands, and receptors annotated. Genes were identified via *DESeq2* (Wald-test) on pseudobulk counts aggregated by biological replicate with *p*-value adjustment using Benjamini-Hochberg method. Genes with |log_2_FC| > 1 and *q* < 0.01 were annotated as significant.

**Extended Data Fig. 4:**
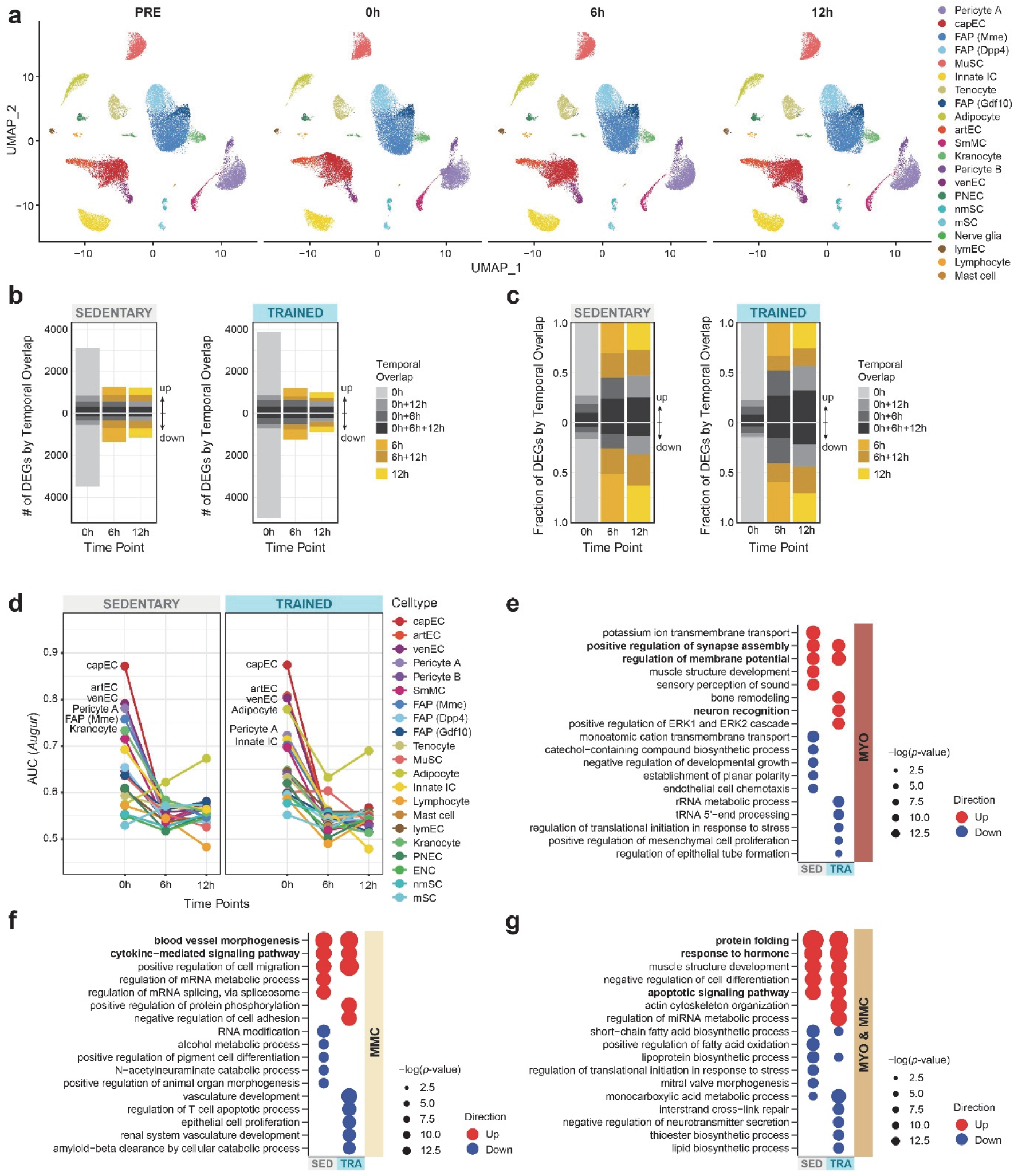
Further analysis of exercise-induced gene expression dynamics. **a**, UMAP projections of MMC fraction from sedentary and trained mice across four time points (PRE, 0h, 6h, 12h). **b**,**c**, Stacked bar plots showing the temporal overlap of DEGs across time points, separated by training state, with absolute DEG numbers (**b**) and relative proportions (**c**). DEGs from all clusters are merged per time point. **d**, Line plots showing *Augur*-inferred perturbation scoring (AUC) of MMC fraction across time points and training states. **e-g**, Gene Ontology enrichment of DEGs in myonuclear (**e**), MMC (**f**), and shared fraction (**g**). Dot size indicates –log10(*p*-value); color denotes directionality (red: upregulated, blue: downregulated). Top five significant terms of each training state and directionality are displayed. SED, Sedentary; TRA, Trained.

**Extended Data Fig. 5:**
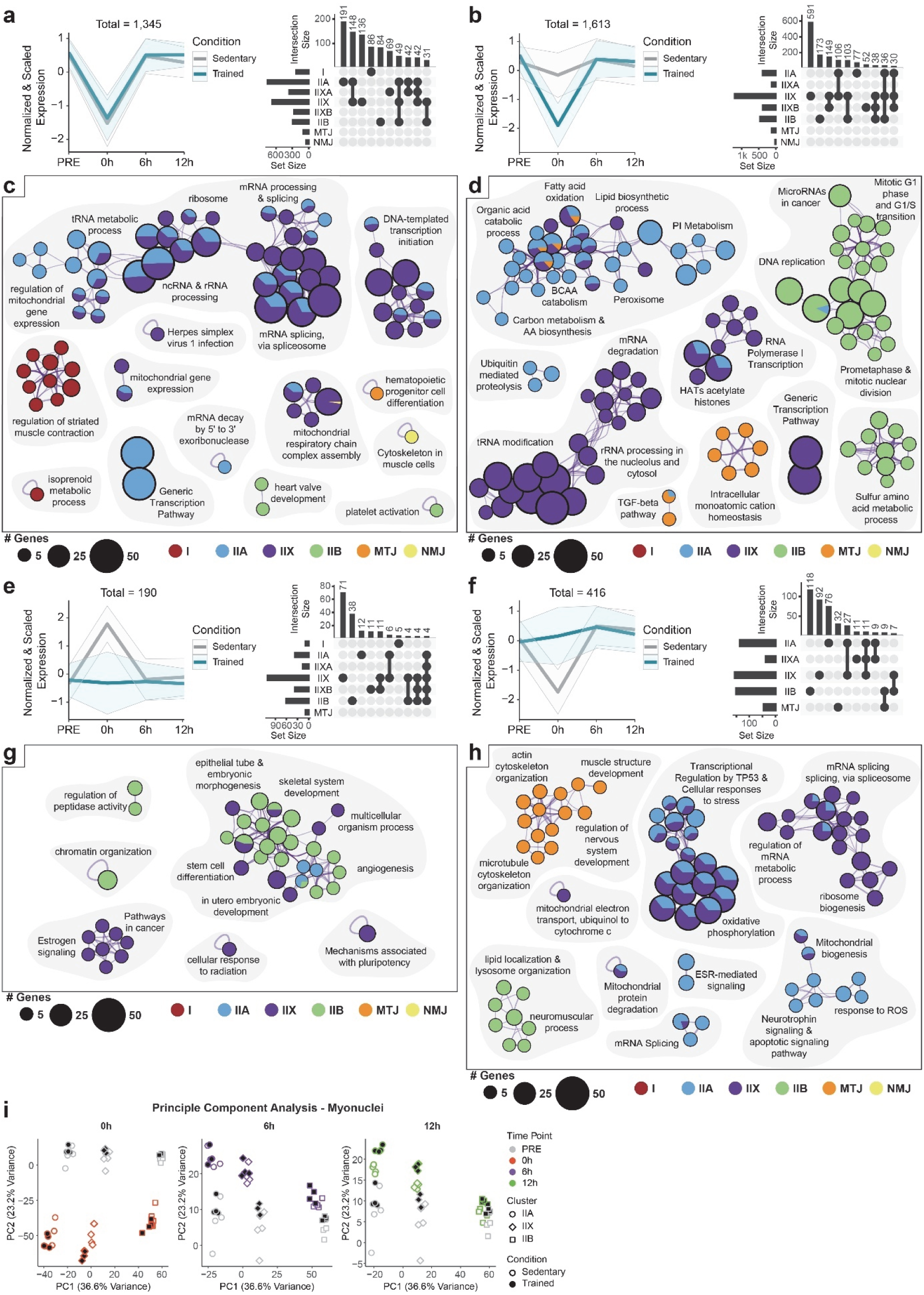
Additional transcriptional modules reveal expanded functional rewiring by training. **a,** Line plot showing normalized and scaled expression patterns of training-state-independently downregulated genes at 0h post-exercise (left) with respective UpSet plot showing contribution of individual myonuclear subtypes (right). **b,** As in **a** but for genes with downregulation in trained muscle. **c**,**d**, Gene Ontology enrichment networks for training state-independently downregulated gene module (**c**) and training-suppressed gene module (**d**). Node size reflects gene set size; node color shows contributing myonuclear subtypes. **e**,**f**, As in **a** and **b** but for genes showing upregulation (**e**) or downregulation (**f**) only in sedentary muscle at the 0h time point. **g**,**h**, Enrichment networks for gene modules in (**e**) and (**f**). **i**, Principal component analysis (PCA) of pseudobulk myonuclear transcriptomes at each time point. Each point represents a biological replicate colored by time and shaped by myonuclear subtype. Fill indicates training state. In **a**,**b**,**e**,**f** curves represent mean expression across myonuclear subtypes ±sd.

**Extended Data Fig. 6:**
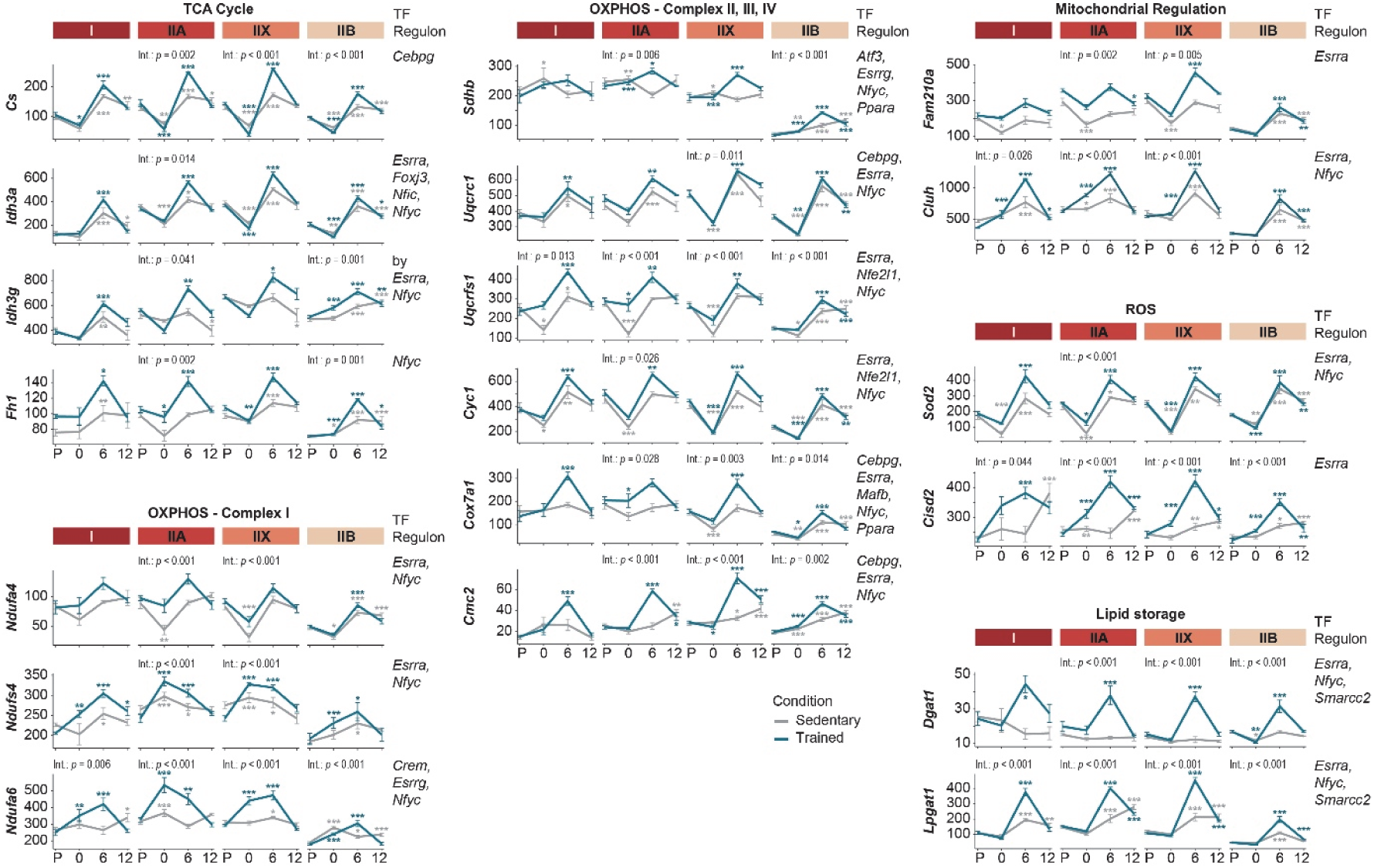
Transcriptional profiles of oxidative metabolism linked to contextual TF networks. Line plot showing time and condition-dependent CPM expression profiles of genes related to oxidative metabolism across main fiber types. Transcription factor names on the right indicate regulons. Data are mean ±sem from pseudobulk profiles (n = 4). Statistical significance assessed by *DESeq2* (Wald-test) comparing time point expression profiles to training state-specific control; *p-*values adjusted with the Benjamini–Hochberg method; \**q* < 0.05, \*\**q* < 0.01, \*\*\**q* < 0.001. Overall interaction via likelihood ratio test. TF: transcription factor.

**Extended Data Fig. 7:**
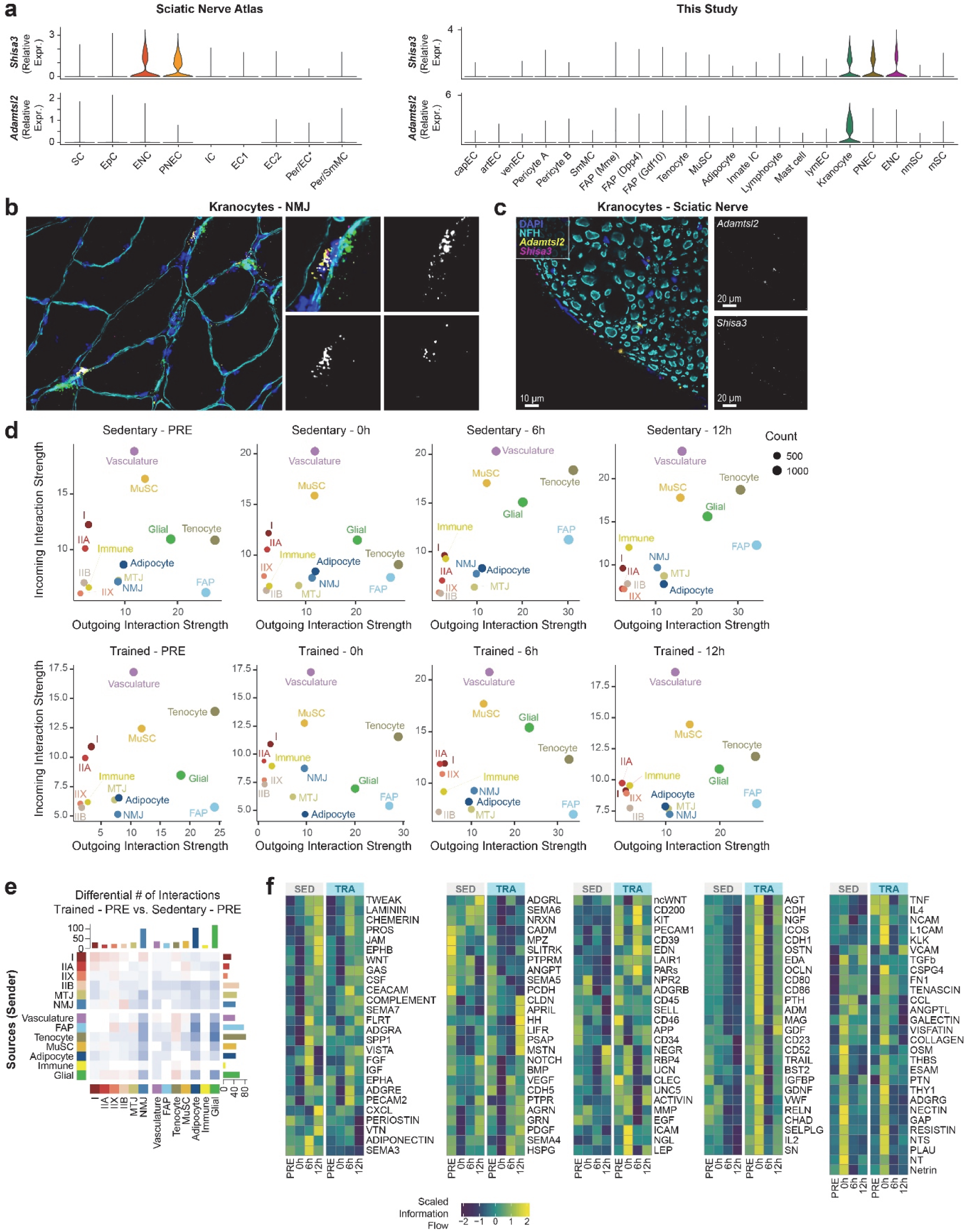
Identification of kranocytes and extended analysis of exercise-induced intercellular signaling. **a**, Violin plots showing *Adamtsl2* and *Shisa3* expression in the sciatic nerve atlas (left) and in the current study (right). Kranocytes (*Mme*⁺ FAP subset) specifically express both markers and are distinct from canonical nerve-associated glia. SC, Satellite cell (neuronal); EpC, Epithelial cell; Per, Pericyte. **b**, Representative RNA FISH images showing colocalization of *Adamtsl2* and *Shisa3* transcripts in kranocytes localized at neuromuscular junctions (NMJs). Inset shows high-magnification views of signal specificity. Blue = DAPI; cyan = Neurofilament H (NFH) for nerve; yellow: *Adamtsl2* transcript; red = *Shisa3* transcript, green = *Chrne* transcript marking the NMJ region. **c**, RNA FISH images from sciatic nerve sections. **d**, Scatter plots showing incoming and outgoing intercellular signaling strength across cell types as predicted by *CellChat*, for sedentary (top row) and trained (bottom row) mice at all time points (PRE, 0h, 6h, 12h). Point size reflects the number of inferred ligand–receptor events per cell type. MMC populations (e.g., FAPs, glial cells, vasculature) exhibit the highest levels of outgoing and incoming signaling. **e**, Heatmap of differential ligand–receptor interaction numbers between trained and sedentary PRE conditions, stratified by sender–receiver cell type combinations. **f**, Scaled pathway information flow across selected *CellChat* signaling categories, showing temporal and training-dependent remodeling of signaling dynamics. Color intensity reflects inferred information flow magnitude (z-score values per row). SED, Sedentary; TRA, Trained.

**Extended Data Fig. 8:**
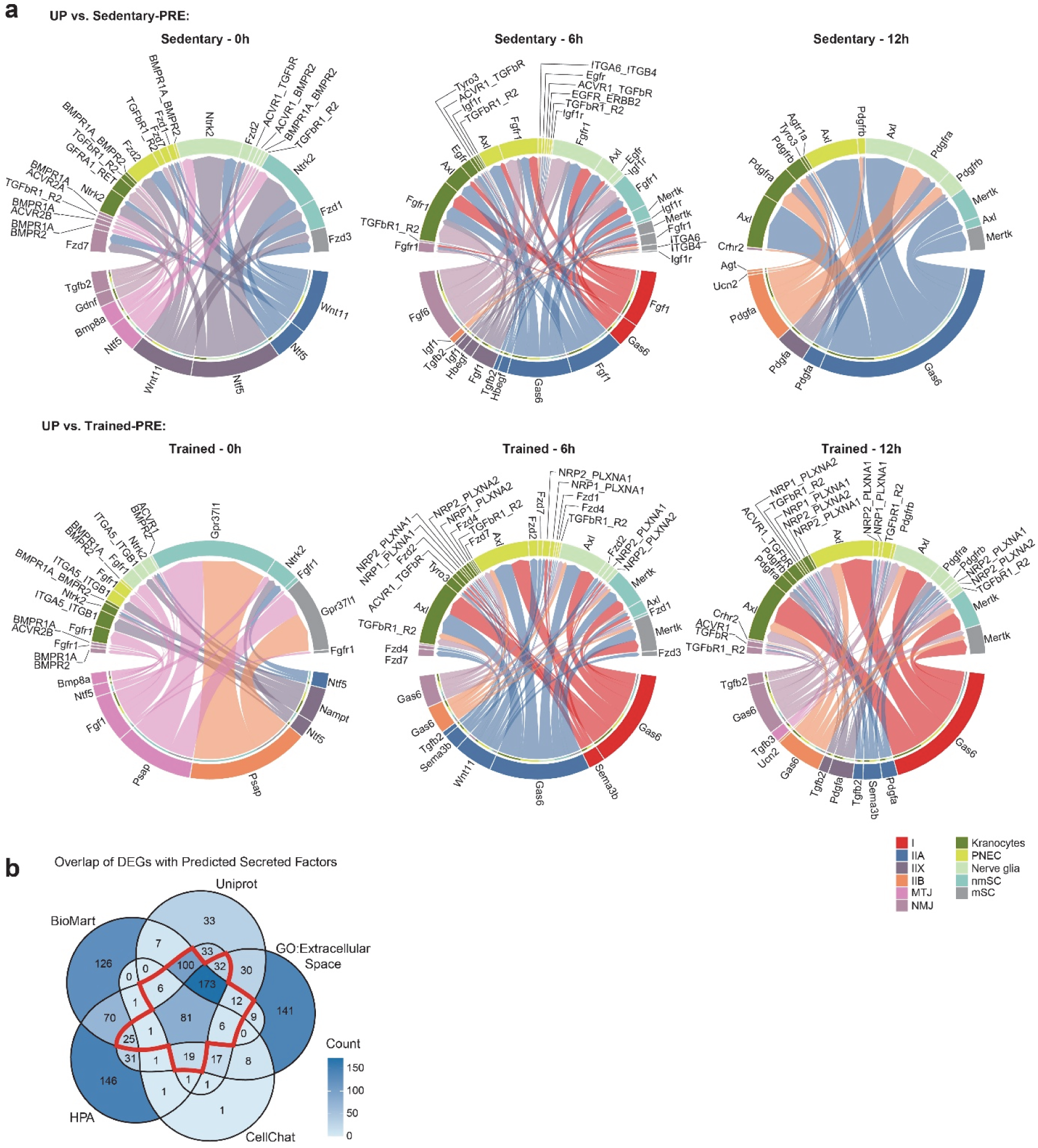
Exercise-dependent ligand-receptor signaling from myonuclei to glia. **a**, Circular chord plots visualizing differentially expressed ligand–receptor pairs between myonuclei and glial subpopulations at 0h, 6h, and 12h following exhaustive exercise compared to the respective PRE control of sedentary and trained conditions. Genes were filtered for log₂FC > 0.5 and *q* < 0.01 for either ligand or receptor. Ligands are shown on the bottom, receptors on the upper side. Colors indicate sender/receiver identity. **b**, Overlap of all DEGs with predicted secreted proteins from external databases. Secretome prediction was based on union across UniProtkb, BioMart, GO:Extracellular space, the Human Protein Atlas (HPA), and *CellChat*’s built-in receptor–ligand database. Red line shows all genes with annotation in at least three of the four databases (independent of *CellChat*) for subsequent GO analysis.

**Extended Data Fig. 9:**
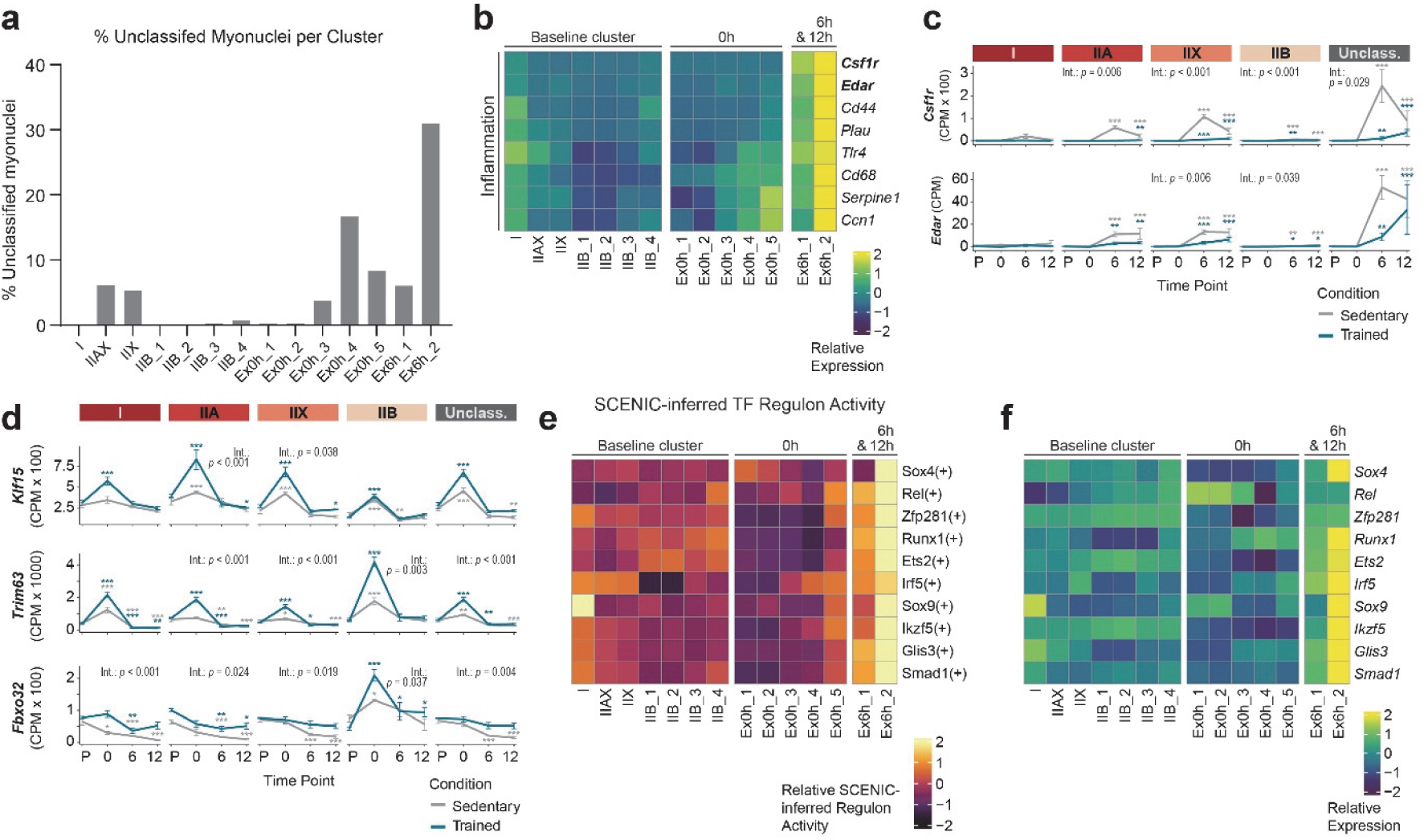
Extended characterization of late-responding myonuclei following exercise. **a**, Bar plot showing the proportion of unclassified myonuclei across all identified myonuclear clusters. **b**, Heatmap showing the relative expression of inflammatory mediators across myonuclear clusters. **c**, Temporal expression profiles of selected inflammatory genes *Csf1r* and *Edar* across myonuclear subtypes. **d**, Expression trajectories of canonical atrophy-related genes (*Klf15*, *Trim63*, *Fbxo32*) across myonuclear subtypes. **e**, SCENIC-inferred transcription factor activity (AUC scores) for the top regulons in *Ex6h_2*. **f**, Heatmap showing relative expression of transcription factors inferred to be active in *Ex6h_2* via SCENIC analysis. In **b**,**e**,**f** pseudobulk aggregated counts or SCENIC-inferred AUC values were normalized for cluster size (CPM) and scaled per row (z-score). In **c**,**d** data represent pseudobulk expression (CPM mean ±sem, n = 4 mice per group). Statistical significance was assessed using *DESeq2* (Wald-test); Benjamini–Hochberg adjusted *p*-values are shown: \**q* < 0.05, \*\**q* < 0.01, \*\*\**q* < 0.001.

**Extended Data Fig. 10:**
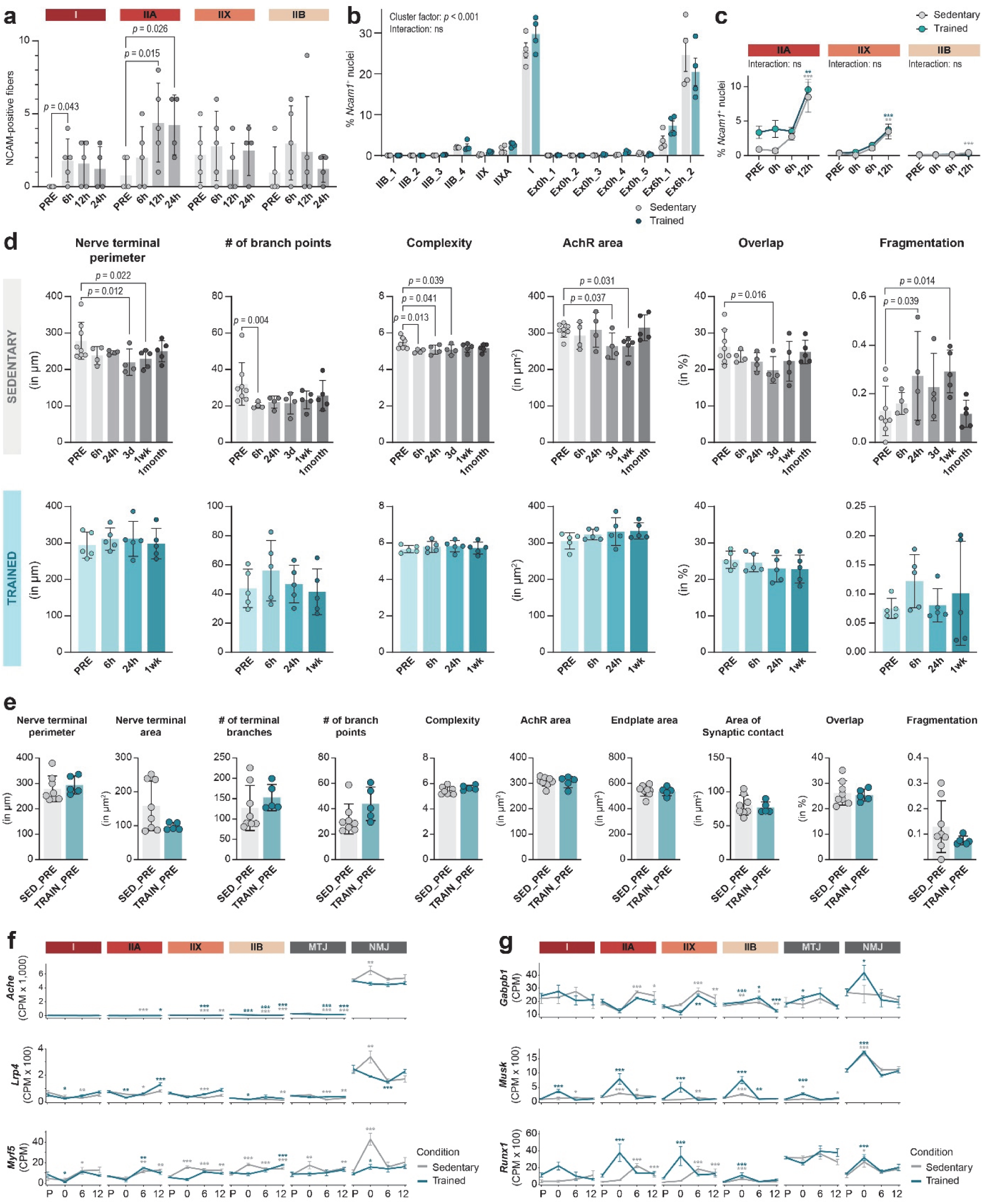
Additional analysis of neuromuscular junction morphology and baseline innervation. **a**, Quantification of NCAM-positive fibers by fiber type, based on immunofluorescent staining (n = 4-5 mice per group). **b**, Percentage of *Ncam1*-positive myonuclei in each population, based on snRNA-seq data. Two-way ANOVA with Tukey’s post hoc test, n = 4 mice per group. **c**, Percentage of *Ncam1*-positive myonuclei in IIA, IIX, and IIB myonuclei after exhaustive exercise in sedentary and trained mice. Two-way ANOVA with Tukey’s post hoc test, n = 4 mice per group, adjusted p-values are shown: \**p* < 0.05, \*\**p* < 0.01, \*\*\**p* < 0.001. **d**, Quantification of NMJ staining using aNMJmorph analysis showing nerve terminal perimeter, number of branch points, complexity, AchR area, overlap and fragmentation in sedentary mice and trained mice upon exercise. **e**, Same as in **d**, showing baseline comparison of sedentary and trained mice. **f**, Line plots showing training-suppressed expression pattern of canonical NMJ genes in sedentary and trained mice post-exercise across myonuclear populations. **g**, Line plots showing training-enhanced expression pattern of NMJ-supporting genes in sedentary and trained mice post-exercise across myonuclear populations. Data are presented as mean ± SD for panel **a**, **d** and **e**. One-way ANOVA with Fisher’s LSD post hoc test was used. When data did not pass normality using by the Shapiro–Wilk test, a non-parametric Kruskal–Wallis test with uncorrected Dunn’s post hoc test was applied. An unpaired t-test was used for baseline comparisons. For panel **d** and **e**, maximum-intensity projections of an average of 23–24 NMJs per mouse were quantified (n = 4–5 mice/group, except for SED_PRE n = 8 mice). In **f**,**g** data represent pseudobulk expression (CPM means ±sem, n = 4 mice per group). Statistical significance was assessed using *DESeq2* (Wald-test); Benjamini–Hochberg adjusted *p*-values are shown: \**q* < 0.05, \*\**q* < 0.01, \*\*\**q* < 0.001.

